# A new evolution-based genomic prediction model forecasts yield performance across environments and future climates and identifies adapted maize landraces

**DOI:** 10.64898/2026.01.28.699851

**Authors:** Agustin O. Galaretto, Marie Pégard, Rosana Malvar, Laurence Moreau, Ana Butrón, Pedro Revilla, Delphine Madur, Valérie Combes, Carlotta Balconi, Cyril Bauland, Pedro Mendes-Moreira, Hrvoje Šarčević, Ana M. Barata, Danela Murariu, Beate Schierscher-Viret, Alex Stringens, Violeta Andjelkovic, Sandra Goritschnig, Brigitte Gouesnard, Alain Charcosset, Stéphane D. Nicolas

## Abstract

Forecasting vulnerability of cultivated and wild species to climate changes is highly challenging. Evolutionary genomic models enable the prediction of mal-adaptation (genomic offset - GO) across environments and future climates under the assumption that populations are currently locally adapted but do not predict but the resulting phenotypic changes. To do so, we developed a new genomic prediction model (GP) integrating both genomic offset (GO) and within-population gene diversity (Hs) to capture genotype by environment interaction and inbreeding effects, respectively (GP-HO-Hs). As proof of concept, we applied this GP-GO-Hs model to a collection of 397 maize populations (landraces) evaluated across 25 environments in Europe using high-throughput DNA pool genotyping. GP-GO-Hs model accurately predicted yield, plant height and flowering time. It increased by 13% the predictive abilities of GP model for predicting yield of new landraces in new environments. GP-GO-Hs model also predicted that the more diverse the landrace, the more stable its agronomic performance across environments. GP-GO-Hs model generated phenotypic adaptive landscapes for each landrace in future climatic scenarios, enabling the identification of landraces with enhanced potential to adapt to future or emerging cultivation conditions. This GP-GO-Hs model could be easily applied to other wild and cultivated species.

**Teaser:** Identify promising landrace adapted to new and future environments by combining genomic selection and offset

## Introduction

Adaptation of crops to changing growing conditions is critical to meet future food demands, as major crops are expected to be negatively impacted by climate change and variability (Ceccarelli et al., 2010; Challinor et al., 2014; Godfray et al., 2010; Minoli et al., 2022). This impact is expected to be variable across regions and crops (Jägermeyr et al., 2021; Rezaei et al., 2023). Maize is the world’s most produced cereal and the one expected to be most highly demanded in the future for food, feed and fuel (OECD & Food and Agriculture Organization of the United Nations, 2022). Maize global production is expected to be negatively impacted by climate change with up to 11% of yield-loss expected in Europe (Heino et al., 2023; Jägermeyr et al., 2021; Knox et al., 2016; Zhao et al., 2017). Breeding for or introducing cultivars adapted to these future conditions could mitigate or even reverse this impact (Challinor et al., 2016; Minoli et al., 2022; Parent et al., 2018; Tsubo & Moeletsi, 2025). Therefore, there is an urgent need for characterizing the available genetic diversity to facilitate its utilization (Martini et al., 2021; Mascher et al., 2019; *FAO*, 2025).

Maize was domesticated in Mexico approximately 9000 years ago and adapted to wide uses and pedo-climatic conditions throughout America, Europe and the rest of the world leading to locally adapted open pollinated populations–landraces (Brandenburg et al., 2017; Matsuoka et al., 2002; Mir et al., 2013; Romero Navarro et al., 2017). Maize landraces have been cultivated in conditions ranging from 0 to 2900 m of altitude, 426 to 4245 mm of annual rainfall and 11.3 to 26.6ºC of annual temperature and in latitudes from 58º N to 40º S (Hallauer et al., 2010; Ruiz Corral et al., 2008). After 1930s, modern hybrids replaced land-races in most parts of the world (Duvick, 2001). But few landraces have been used as the starting point of modern breeding programs and the pedigree of currently cultivated hybrids traces back to only a few founder lines (Arca et al., 2023; Smith et al., 2022).

Despite major genetic progress achieved for yield and abiotic stress tolerance, the narrow diversity of elite germplasm can jeopardize future genetic gains (Allier et al., 2019; Campos et al., 2006; Sanchez et al., 2023; Smith et al., 2022; Welcker et al., 2022) and cope with new traits required for sustainable agriculture (e.g., reduced nitrogen fertilization) or climate change (e.g., increasing temperatures and more frequent droughts) (Hawkesford & Griffiths, 2019; Heino et al., 2023). Consequently, maize landraces are receiving increasing attention as a reservoir of favorable alleles to introduce useful diversity into breeding programs (Böhm et al., 2017; Gorjanc et al., 2016; Hoisington et al., 1999; Hölker et al., 2019; Mayer et al., 2020). However, the utilization of genetic resources is partially constrained by the lack of information, as only 22% of maize accessions have been characterized despite big efforts made by genebanks (*FAO*, 2025). Because the identification of promising accessions is crucial for pre-breeding programs, methods that allow one to predict performances of not yet phenotyped accessions, such as genomic prediction (GP), could have a major impact in breeding for adaptation (Böhm et al., 2017; El Hanafi et al., 2023; Mascher et al., 2019).

GP enables the estimation of breeding values for accessions without phenotypic records using molecular markers and it has long been proposed as a solution to avoid the evaluation of large number of genotypes (Bernardo, 1994; Meuwissen et al., 2001). GP has been widely adopted in maize breeding programs but its application to maize landraces remains limited (Gholami et al., 2021). Previous works have focused on using single plant genotyping to predict the performance of landraces or their derived doubled haploid lines in combination with testers (Böhm et al., 2017; Hölker et al., 2019, 2022; Li et al., 2025; Romero Navarro et al., 2017). These studies have yielded promising results to identify beneficial alleles and represent an important step towards integrating landraces into breeding programs. However, the use of tester may blur the adaptation signal of landraces (Li et al., 2025) and generating DH lines from landraces is a costly process that allows to explore only few selected landraces (Böhm et al., 2017). Thus, efficient GP approach that enable the prediction of landrace *per se* agronomic performance and adaptation in various environments and the identification of best landraces are needed. In addition, genotyping single plants is insufficient to represent within landrace diversity. This is because maize landraces are heterogeneous populations, thus sampling multiple individuals is necessary to accurately characterize each accession, leading to increased genotyping costs.

To cope with this issue, high-throughput pool genotyping (HPG) methods, such as DNA-pool genotyping or sequencing, offer a cost-effective alternative to individual genotyping and could therefore enable the large-scale application of GP to maize landraces (Arca et al., 2021). In forage crops, HPG-based genomic prediction has proven useful for predicting the value of synthetics and natural populations (Cericola et al., 2018; Keep et al., 2020; Pégard et al., 2023). DNA-pool genotyping enabled a detailed understanding of maize landraces’ genetic composition, the history of maize dispersion, within-population gene diversity, and the mechanisms underlying adaptation to diverse environments (Arca et al., 2023; Balconi et al., 2024; Mir et al., 2013; Rebourg et al., 2003; Tamang et al., 2024). DNA-pooling genotyping also showed that within-population gene diversity of maize landraces greatly varied and was positively correlated with their *per se* agronomic performance reflecting certainly inbreeding depression effect (Rebourg et al., 2001). Despite previous work, HPG has not been explored yet to perform GP for maize landraces *per se*.

In addition to GP, ecology-derived methods that predict (mal)adaptation to present or future scenario climatic conditions are gaining attention and have been applied to identify useful genetic resources (Fitzpatrick et al., 2025; Rellstab et al., 2021). These methods link current patterns of adaptive genomic composition to climatic conditions. Ultimately, they allow one to estimate the allelic frequencies shift required at certain loci for populations to be adapted to a new environment, a concept known as genomic offset (GO) (Fitzpatrick & Keller, 2015). Gradient Forest (GF) is one of the most used models to predict (mal)adaptation (Capblancq et al., 2020; Ellis et al., 2012). For maize landraces, GF was used to assess future (mal)adaptation for a set of Mexican and Himalayan landraces, raising concern about their *in-situ* preservation (Aguirre-Liguori et al., 2019; Tamang et al., 2024). Despite the potential of GO to identify endangered or interesting genetic resources for substitution or pre-breeding, the integration of the GO with GP to predict *per-se* maize landrace performances remains unexplored (McLaughlin et al., 2024).

In this study, we investigated the use of GP in combination with GO as a tool to unlock genetic diversity in gene banks by massively predicting maize landrace performances, particularly for complex traits of agronomic and economic importance such as grain yield, flowering time and plant height in various environments and future climatic scenarios. We combined genotypic data obtained from cost-effective HPG, with the phenotypic performances for three agronomic traits of 397 landraces evaluated across a sparse 25 environment network within the ECPGR EVA Maize Network. First, we took advantage from the climatic characterization of both landrace collection sites and field experiments to predict landrace (mal)adaptation to each experiment using GO and how it was related to agronomic performance. Second, we evaluated accuracy of GP based on HPG for male flowering time, grain yield and plant height in three scenarios of increasing difficulty: (i) completing missing data in the trial network, (ii) predicting the values of a new accession in different environments within the network (iii) and predicting a new accession in a new environment. Third, we assessed to which extent within-population gene diversity (Hs) and GO captured part of the genotypic and the genotype-by-environment variation and their potential to improve GP accuracy in these three different scenarios. Finally, we proposed an approach that combined GP and eco-genetic statistics to identify promising accessions for adaptation to future climate scenarios across Europe.

## Results

### Modelling landrace adaptation to past and future climates

To investigate the adaptation of landraces to local environments, we applied the Gradient Forest (GF) approach to a panel of 470 European maize landraces (**Table S1**), combining allelic frequency and climatic variables of landrace collection sites (**Table S2**). GF models allelic frequency variation of loci along climatic gradients. Our GF model identified 243 SNPs significantly associated with the variation of 55 climatic variables from WorldClim2.1 (*R*^2^ > 0.47 –**Figure S1A**). Among these SNPs putatively involved in climatic response, we identified *loci* known to be involved in flowering time and adaptation in previous studies such as *Vgt2* (PZE-108070380; *R*^2^ = 0.60), *Vgt3* (PZE-103098779; *R*^2^ = 0.49) and *ZCN2* (PZE-104057075; *R*^2^ = 0.47, **Table S3**). The GF model ranked the importance of climatic variables to predict allelic frequency variations of these 243 SNPs. This enabled us to retain the top 15 complementary climatic variables (accuracy importance > 0.01; **Figure S1B, Table S2)**.

We then studied spatial genetic variation based on the GF model adjusted with the selected 243 SNPs and these 15 climatic variables (**Figure 1A**). The geographical distribution of genetic composition predicted by GF differentiated several climatic regions. “Green” regions represented the largest area in Europe and included all Eastern European Countries but also Po Valley in North of Italy, Central Southern Spain and Turkey. “Orange” regions were mainly located along the Atlantic Ocean but also along the Mediterranean coast and southern regions of Portugal, France and Italy, with a South-North gradient corresponding to decreasing temperature in summer (light to dark from South to the North). “Pink” regions included Galicia, Northern Western Spain and South Western France. “Blue regions” were the smallest area including mainly mountain regions (Alpes, Massif Central and Pyrenean) as well as German regions. The biplot (**Figure 1B**) presents the grids (∼4.5 x 4.5 km) of the geographical map with their corresponding color as well as the contribution of each variables to PCA axis. In Figure 1B, the first PCA axis, linked with red color intensity, was associated with winter temperature (*tmax_11*), while the second axis, linked with green color intensity, was associated with temperature seasonality (*bio_4*) and temperature annual range (*bio_7*). The first axis discriminated landrace groups of the Iberian regions (SE-Spanish, Galician and Portuguese landraces), where winter temperature is soften by the oceanic effect, from landraces assigned to the Northern Flint (NF) and Corn Belt Dent (CBD) genetic groups (Balconi et al., 2024). The second axis discriminated landraces assigned to the CBD group, collected in continental regions where temperature varies greatly across the year, from NF and Galician groups. The geographic distribution of genetic groups strongly mirrored the genetic composition predicted by GF, suggesting local adaptation of genetic groups.

**Fig. 1.**
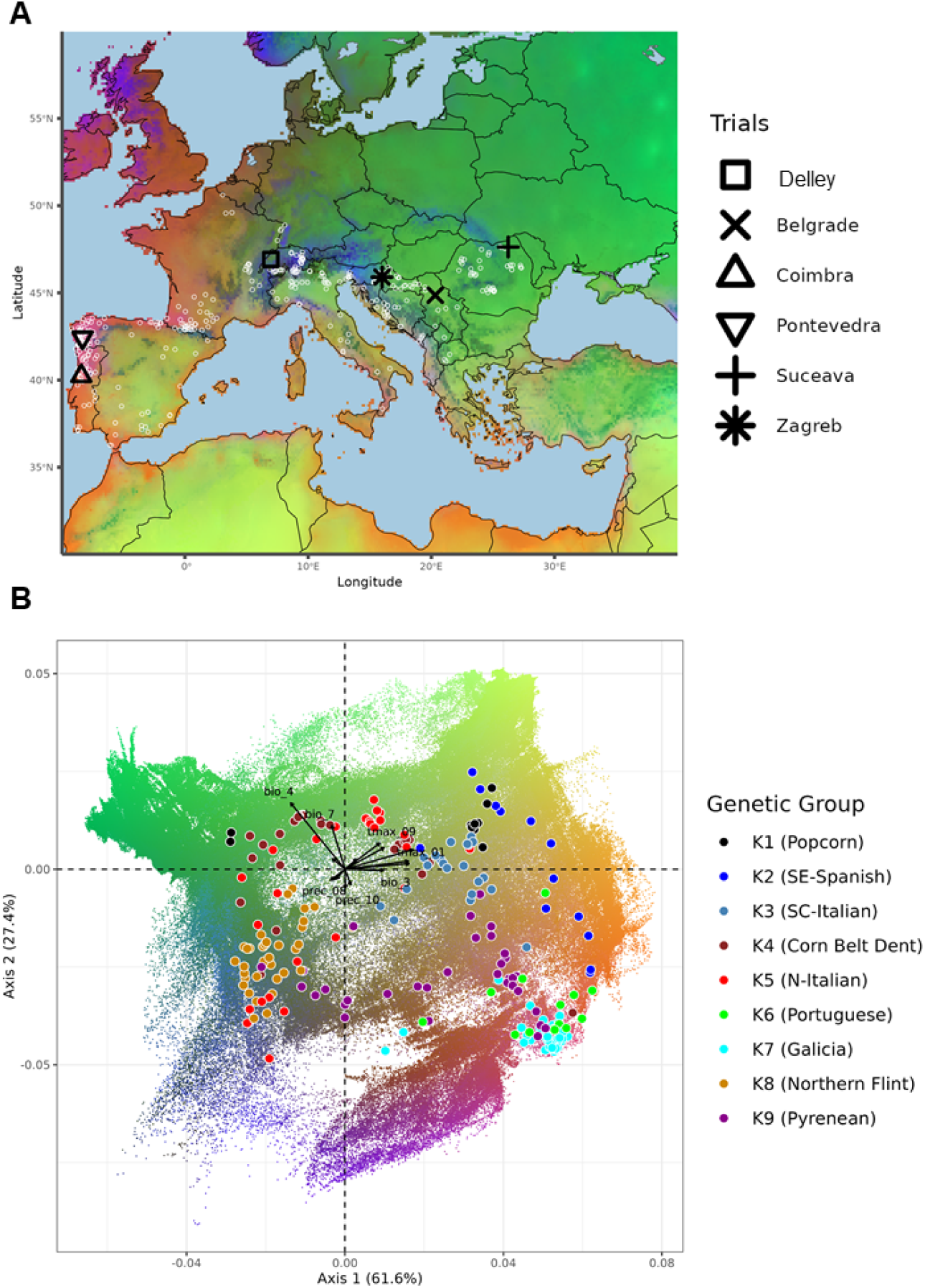
Genetic adaptive landscape of maize landraces to past climatic conditions (1970-2000). **(A)** Adaptative landscape map of maize landraces is colored according to the first three axes of the Principal Component Analysis (PCA) on Gradient Forest (GF) transformed climatic variables from Worldclim 2.1 for period 1970-2000. The color at each grid (∼4.5km^2^) of the geographical map is based on a multivariate scale where each color is a mixture of red, green and blue with its intensity proportional to the first three PCA coordinates, respectively. As GF-transformed climatic variables represent adaptive allelic frequency variation, the different colors illustrate different adaptive genetic composition. Landrace collection sites and EVA Network field experiment sites are represented by white circles and black symbols, respectively. **(B)** The biplot of the first plan of PCA on GF-transformed climatic variables for the grids in the geographical map with their corresponding color and climatic variable vectors. Black symbols represent the projection of climatic conditions at field experiments. Colored circles represent the projection of collection site climatic conditions for landraces highly assigned to genetic groups (> 0.6).

We used the GF model to estimate GO expected under future climatic scenarios for each grid of the map across Europe (local GO). To do so, we used GF-transformed climatic variables from WorldClim2.1 past (1970-2000) data and four future climatic scenarios derived from two CO_2_ concentration ([CO2]) corresponding to two greenhouse gases emission trajectories (one optimistic -SSP126- and one pessimistic -SSP585) and two time periods (one at short term - 2041-2060 - and one at long term 2080-2100) with 13 global circulation models (GCMs, **Table S4**). Mean local GO across 13 GCMs increased across Europe under all future climatic scenarios with the highest increase for SSP585 scenario in 2081-2100 horizon, in NE-France, Germany and E-Europe (**Figure S2**). This implies that landraces collected in these regions could be maladapted to future climatic conditions in these same regions.

Similarly, we used the GF model to predict the adaptive landscape for each landrace across Europe. The geographic distribution of GO predictions of four landraces representing different genetic groups showed contrasting patterns for past and future scenarios (**Figure 2**). For past climatic data, two landraces (ESP0148 and FRA0296) showed low GO (<0.03) in large geographic regions close to their collection sites, in Northern and Southern Europe, respectively. In contrast, for BIH0063 and FRA0491, geographic areas with low GO were more limited while those with high GO (> 0.07) were larger and included important maize production regions. Future climatic conditions had a negative impact on local adaptation for these 4 landraces irrespective of the period and the scenarios, with the highest negative impact observed for the 2081-2100 time horizon.

**Fig. 2.**
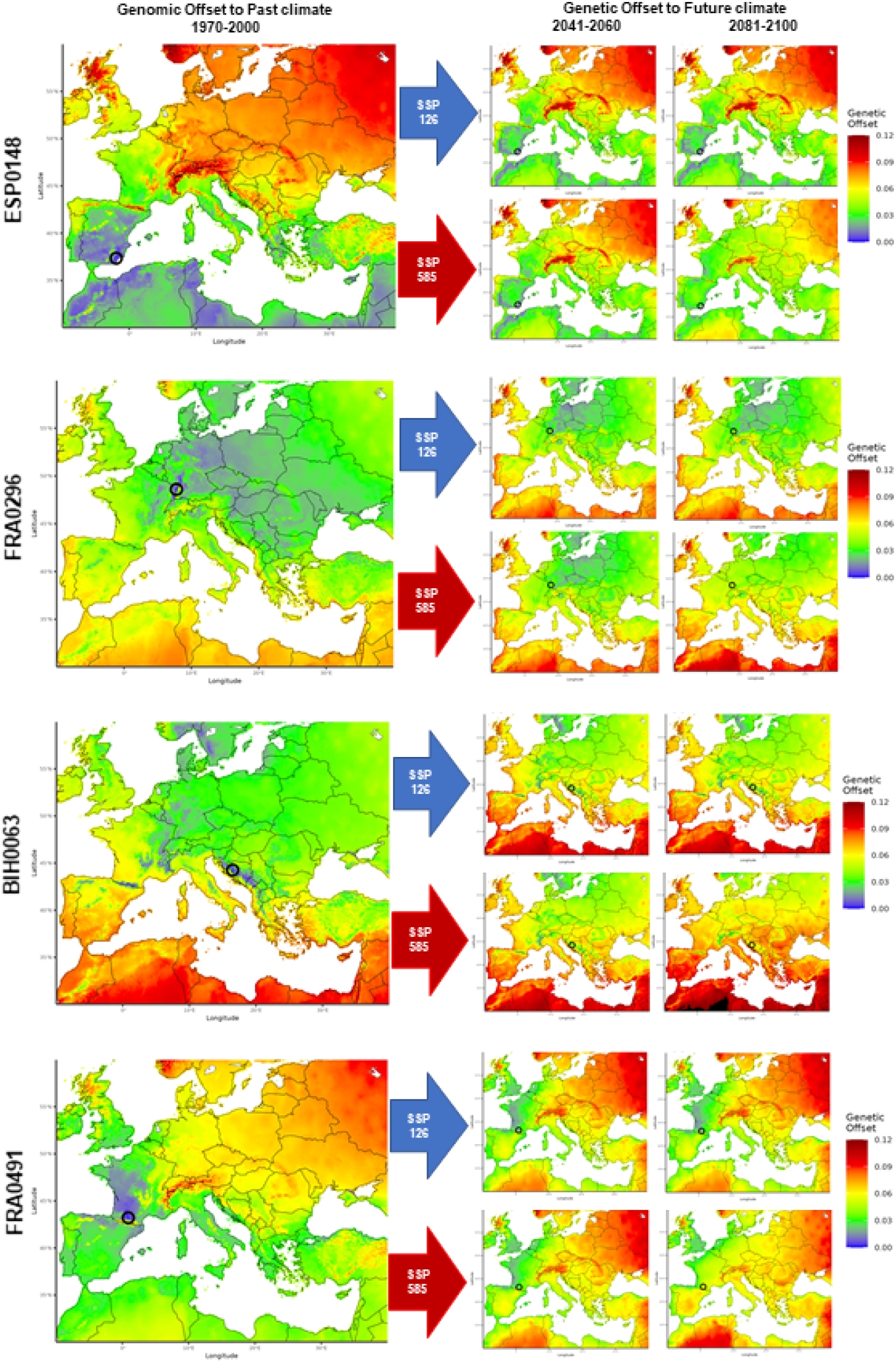
Prediction of (mal)adaptation (genomic offset) in past and future climatic conditions under different emission scenarios. The maps represent GO for four landraces under past (left) and future (right) climatic conditions. GO is defined as the Euclidean distance between GF-transformed past climatic variables of landrace collection sites (black circles) and climatic variables across Europe, in the past (left) or as predicted in the future (right). Because GF models the relationship between climatic variables and adaptive allelic frequencies, this distance represents the genetic distance to a theoretically-local adapted population (i.e adaptation gap). We selected for illustration four landraces representing different genetic groups: ESP0148 (SE-Spanish), FRA0296 (NF), BIH0063 (CBD), and FRA0491 (Pyrenean). Past climatic conditions are based on WorldClim 2.1 (1970-2000) while future climatic scenarios represent the combination of two time-horizons (2041-2060 and 2081-2100) with two emission pathways: SSP126 (blue arrows) and SSP585 (red arrows).

The pessimistic scenario (SSP585) led to an increase of the geographical area with high GO, and a reduction of geographical area with low GO. In contrast, under the optimistic scenario (SSP126), the model showed globally a small decrease in low GO area. The landrace BIH0063, assigned to CBD, was the most affected by climate change. In contrast, the SE-Spanish landrace ESP0148 showed less dramatic changes in GO for future scenarios. As GO relies only on climatic and genotypic data, no inference can be done in terms of loss of agronomic performance, limiting the interpretation of maladaptation predictions based on GO only.

### Effect of genomic offset and within-population gene diversity on agronomic traits

In order to test the effect of (mal)adaptation predicted by GO on agronomic performances, we evaluated the phenotypic response of 397 landraces to 25 environments for male flowering time (MF, in growing degrees units-GDU), plant height (PH, in cm) and grain yield (GY, in t/ha), using a sparse experimental design (**Table S5**). The results of phenotypic evaluation are summarized in **Table 1** and **Table S6**. Means across the network were 788 GDU for MF, 183 cm for PH and 4 t/ha for GY (**Table 1**). Mean heritabilities were 0.96 for MF, 0.91 for PH and 0.86 for GY (**Table 1**). Significant variation of the three traits was observed between and within experiments (**Figure S3, Table S6)**. The maximum GY was achieved at Zagreb (Croatia) in 2024 while the lowest one was recorded at Zagreb (Croatia) in 2022 (**Table S6**).

**Table 1.** Summary of EVA network environment means and heritabilities for male flowering, plant height and grain yield. Mean, standard deviation (SD), minimum and maximum is based on 25 environments of the EVA network.

| Trait | Acronym | Unit | EVA network phenotypic data |  |  | EVA network heritability |  |  |
| --- | --- | --- | --- | --- | --- | --- | --- | --- |
|  |  |  | Mean (SD) | Min | Max | Mean (SD) | Min | Max |
| Male Flowering | MF | GDU | 788.64 (103.8) | 624.84 | 1007.89 | 0.964 (0.03) | 0.9 | 0.99 |
| Plant Height | PH | cm | 186.9 (32.51) | 126.53 | 238.01 | 0.906 (0.09) | 0.64 | 0.99 |
| Grain Yield | GY | t/ha | 3.82 (0.97) | 2.09 | 5.63 | 0.855 (0.12) | 0.44 | 0.99 |

To assess the adaptation of landraces to field experiment conditions, we used GF to estimate GO between environmental conditions of each experiment and each landrace collection site (**Table S7**). Our field trial experimental network covered well the range of climatic conditions of landraces’ collection sites (**Figure 1B**). Maximum GO (corresponding to putatively the worst adapted accession) was 0.107 for the landrace “Vilariño” (ESP0142), collected in Galicia and evaluated at Suceava (Romania) in 2021 (SSR_2021_1A). In contrast, the lowest GO (putatively best adapted accession) was 0.015 for the Portuguese landrace “Milho Amarelo” (PRT0202) evaluated at Pontevedra (Galicia, Spain) in 2021 (CPS_2021_1A). The GO range for each landrace across the field trial experimental network (maximum GO – minimum GO) was highly variable (0.0002 to 0.0875, **Figure S4**), indicating that only part of the landraces (∼10%) was evaluated in contrasted conditions of low (<0.03) and high (>0.07) GO conditions. GO and the geographic distance between landrace collection sites and field experiments were correlated (**Figure S5**), indicating that the lowest GO were reached for landraces collected close to the experimental trials. This further supports the close relationship between local adaptation and the geographic distribution of genetic groups.

To assess the effect of GO estimation of maladaptation on phenotypic variation, we analyzed the relationship between GO and traits measured at each trial 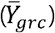. GO was slightly but significantly negatively correlated with GY (*p* < 0.001, *b* = -19 t/ha, *R*^2^ = 0.05; **Figure 3A**) and PH (*p* < 0.001, *b* = -464 cm, *R*^2^ = 0.06; **Figure S6A**). In the contrary, we observed no relationship between GO and MF (*p* = 0.173, *R*^2^ < 0.001; **Figure S6B**). Focusing only on landraces that were evaluated under contrasting GO conditions (*n* = 46) increased the linear relationship between GO and GY (*b* = -27 t/ha, *R*^2^ = 0.16), while this only had a limited or no effect for PH and MF, respectively (**Figure S7**). This suggests that the effect of GO on decreasing agronomic performance was certainly underestimated for the entire panel, since most landraces were evaluated in trials with weak contrasting GO contrast.

**Fig. 3.**
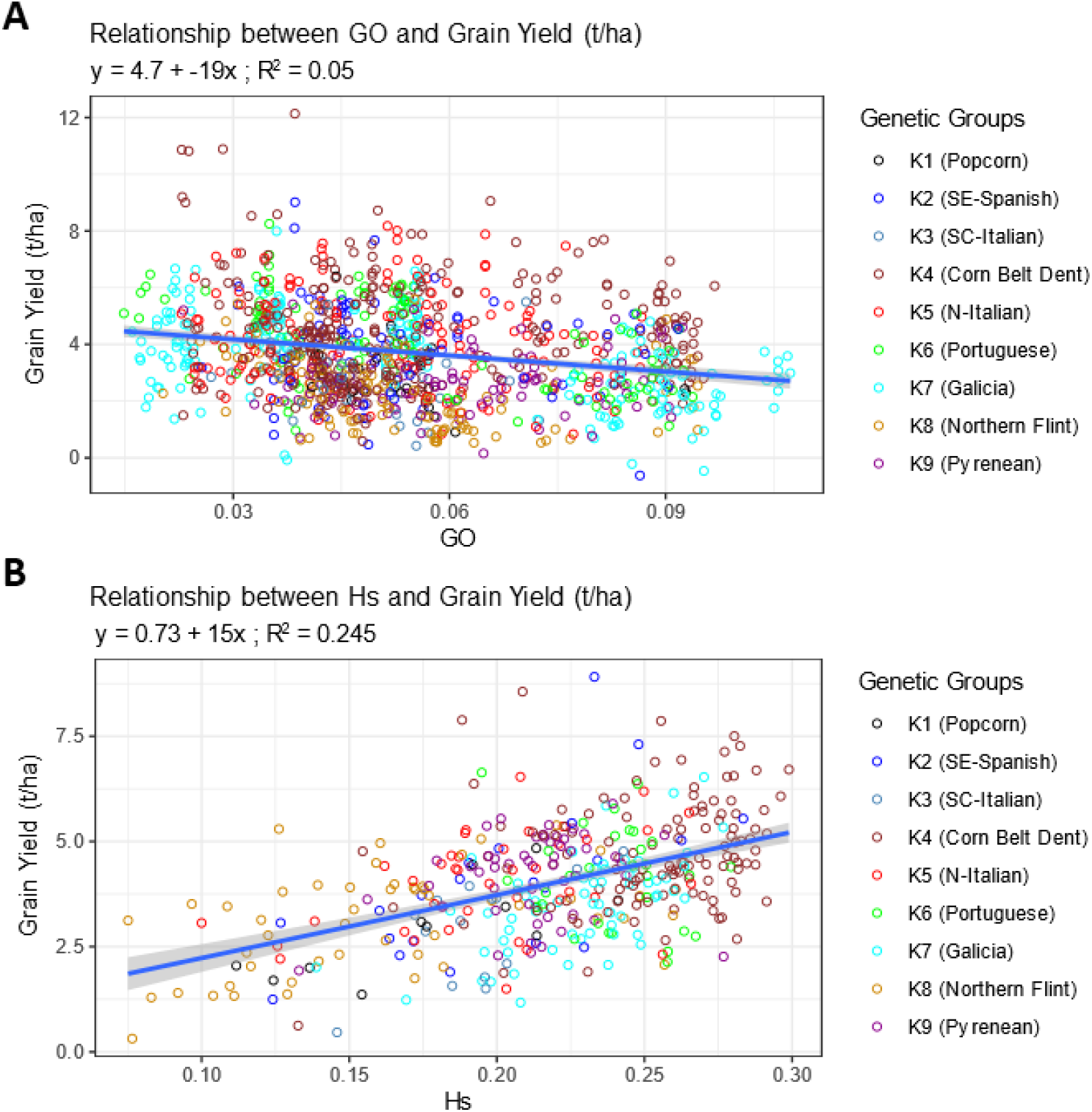
Effect of within-population gene diversity (Hs) and genomic offset (GO) on grain yield (GY). **(A)** Relationship between genomic offset (GO) of each landrace to each field experiment and grain yield within experiment adjusted means. Each circle represents one landrace evaluated in one experiment. **(B)** Relationship between within-population gene diversity (Hs) with grain yield least square means across EVA Network field experiments. Each circle represents one landrace. In **A** and **B**, circles are colored by maximum assignment to 9 genetic groups.

We also assessed the effect of inbreeding depression on phenotypic variation by studying the relationship between within-population gene diversity (Hs) and the three traits (**Figure 3B and Figure S8**). Mean Hs was 0.214 and it varied greatly among landraces, ranging from 0.075 (CHE0355 assigned to NF) to 0.299 (ITA0083 assigned to CBD) (**Table S1)**. Hs showed a significant and positive linear relationship with GY (*p* < 0.001, *b* = 15 t/ha, *R*^2^ = 0.25; **Figure 3B**) and PH (*p* < 0.001, *b* = 321 cm, *R*^*2*^ = 0.30; **Figure S8A**) but not with MF (*p* = 0.558, *R*^*2*^ < 0.01; **Figure S8B**). This suggests that flowering time is less affected by inbreeding depression than GY and PH.

### Within-population gene diversity and genomic offset capture a part of genotypic and genotype-by-environment variance of grain yield and plant height

To evaluate the contribution of Hs and GO to the genotype and genotype by environment (G×E) variance components, we compared a first model (M1) accounting for the genotype and G×E effects with models also including either the GO effect (M2), or the Hs effect (M3) or both (M4). (**Table 2**).

**Table 2.**
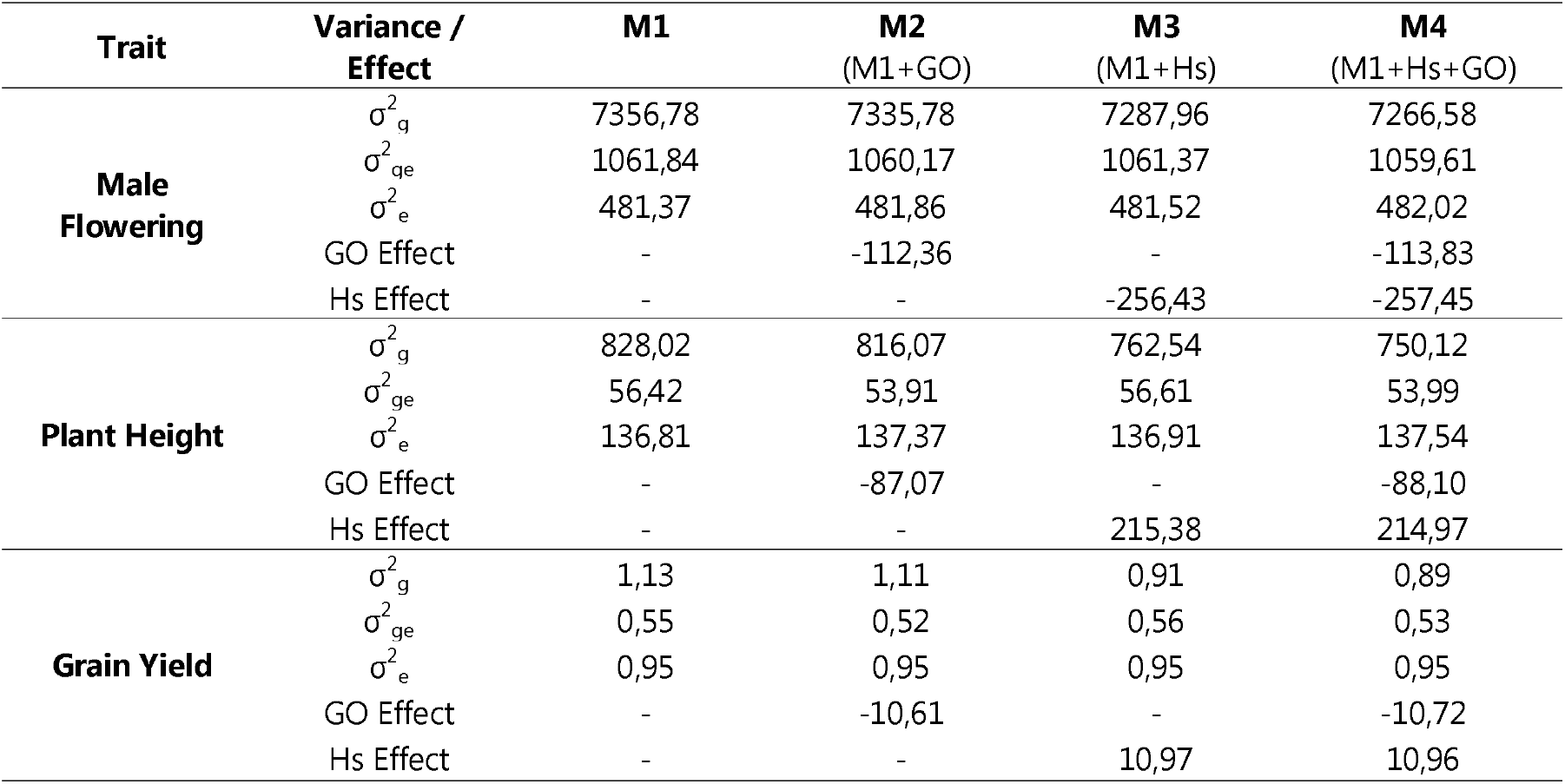
Variance decomposition of male flowering, plant height and grain yield for the four models. Base model M1 included genotype, genotype-environment interaction as random effects. M2 model included all terms of M1 model plus genomic offset (GO) as fixed effect. M3 model included all terms of M1 model plus within-population gene diversity (Hs) as fixed effect. M4 model included all terms of M1 model plus both GO and Hs as fixed effect.

| Trait | Variance / Effect | M1 | M2<br>(M1+GO) | M3<br>(M1+Hs) | M4<br>(M1+Hs+GO) |
| --- | --- | --- | --- | --- | --- |
| Male Flowering | $\sigma_g^2$ | 7356,78 | 7335,78 | 7287,96 | 7266,58 |
| | $\sigma_{ge}^2$ | 1061,84 | 1060,17 | 1061,37 | 1059,61 |
| | $\sigma_e^2$ | 481,37 | 481,86 | 481,52 | 482,02 |
|  | GO Effect | - | -112,36 | - | -113,83 |
|  | Hs Effect | - | - | -256,43 | -257,45 |
| Plant Height | $\sigma_g^2$ | 828,02 | 816,07 | 762,54 | 750,12 |
| | $\sigma_{ge}^2$ | 56,42 | 53,91 | 56,61 | 53,99 |
| | $\sigma_e^2$ | 136,81 | 137,37 | 136,91 | 137,54 |
|  | GO Effect | - | -87,07 | - | -88,10 |
|  | Hs Effect | - | - | 215,38 | 214,97 |
| Grain Yield | $\sigma_g^2$ | 1,13 | 1,11 | 0,91 | 0,89 |
| | $\sigma_{ge}^2$ | 0,55 | 0,52 | 0,56 | 0,53 |
| | $\sigma_e^2$ | 0,95 | 0,95 | 0,95 | 0,95 |
|  | GO Effect | - | -10,61 | - | -10,72 |
|  | Hs Effect | - | - | 10,97 | 10,96 |

For M1, the genotype variance 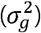 accounted for 83%, 81% and 43% of the total variance for MF, PH and GY, respectively. The G×E variance 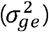 accounted for 12%, 6% and 21% of total variance for MF, PH and GY, respectively. This indicates that G×E contributed approximately two and four times more to GY variation than to MF and PH, respectively. When adding GO in the model (M2), 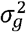 slightly decreased in reference to M1 for PH and GY (-1% and -2%, respectively). This decrease was stronger (-4% and -6%, respectively) for 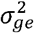, indicating that GO captured mainly G×E for both PH and GY. When Hs was included in the model (M3), 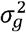 decreased by 8% and 20% for PH and GY relative to M1, respectively, while there was no or a slight increase in 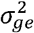. This indicates that Hs captured mainly the genetic variation between landraces for PH and GY. Finally, when including both GO and Hs (M4), we observed a decrease of the genotypic and G×E variance for PH and GY which was much stronger for 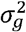 (-9% and -22%, respectively) than for 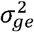 (-4% for both traits). Hs and GO captured therefore 19% and 10% of the genetic variance component for GY and PH, respectively. GO and Hs acted independently on trait variation since Hs and GO displayed similar effects irrespective of whether they were considered together in the model (M4) or as unique factors of variation (M2 and M3). With the model 4, an increase of GO by 0.1 reduced GY by 1.1 t/ha and PH by 88 cm, while an increase of Hs by 0.1 increased GY by 1.1 t/ha and PH by 214 cm (**Table 3**). This suggests that PH is proportionally more impacted by Hs than GO compared to GY (**Table 3**).

**Table 3.**
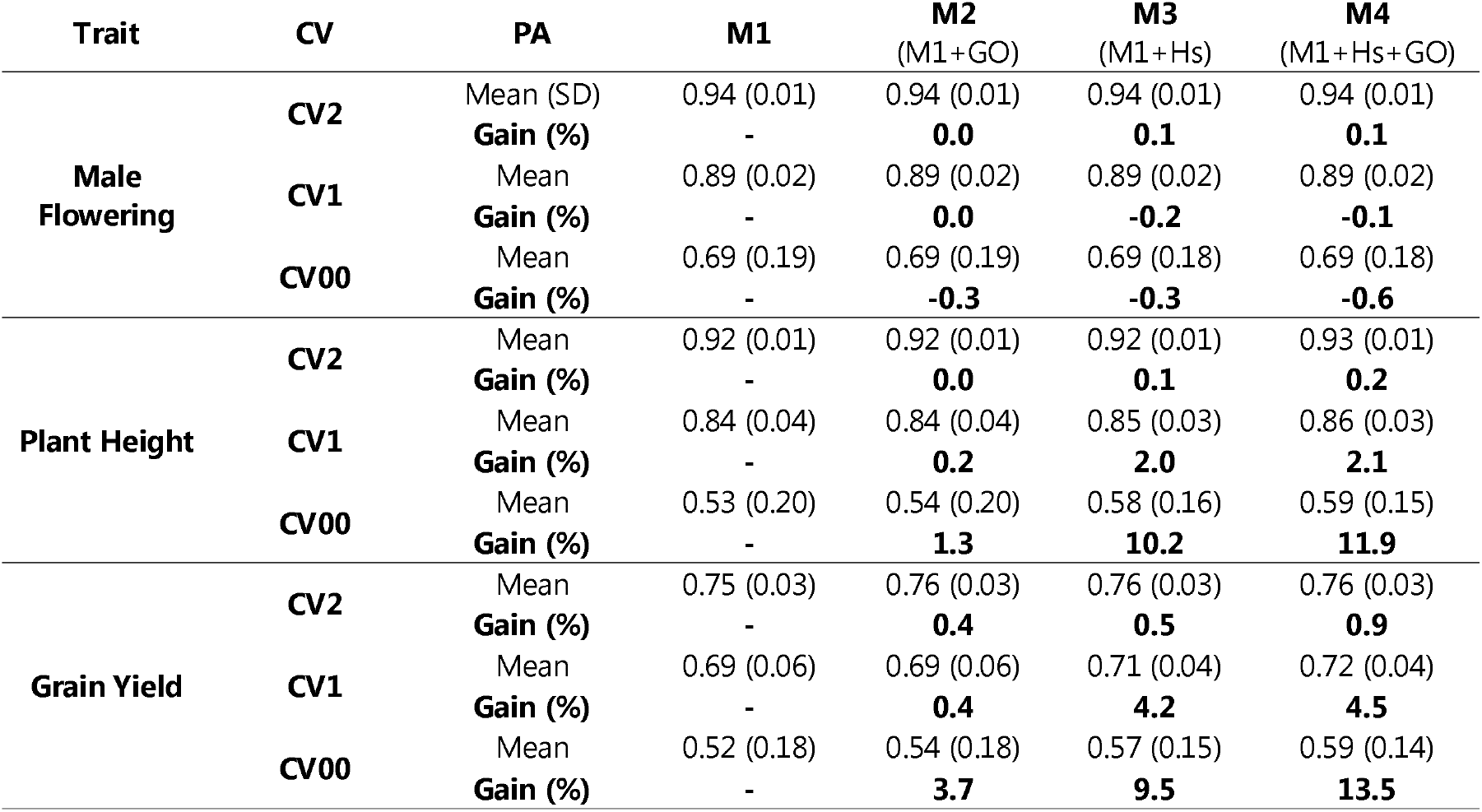
Comparison of predictive abilities between four genomic prediction models for three traits that included or not within-population gene diversity and genomic offset and using different cross-validations schemes. Mean, standard deviation (SD) and gain of predictive abilities (PA) were estimated for the different genomic prediction models for the three traits and the three different cross validation schemes (CV). Four models for genomic prediction corresponded to the same models used in Table S2. Predictive abilities gains are estimated in reference to the model M1.

| Trait | CV | PA | M1 | M2<br>(M1+GO) | M3<br>(M1+Hs) | M4<br>(M1+Hs+GO) |
| --- | --- | --- | --- | --- | --- | --- |
| Male Flowering | CV2 | Mean (SD) | 0.94 (0.01) | 0.94 (0.01) | 0.94 (0.01) | 0.94 (0.01) |
|  |  | Gain (%) | - | **0.0** | **0.1** | **0.1** |
|  | CV1 | Mean | 0.89 (0.02) | 0.89 (0.02) | 0.89 (0.02) | 0.89 (0.02) |
|  |  | Gain (%) | - | **0.0** | **-0.2** | **-0.1** |
|  | CV00 | Mean | 0.69 (0.19) | 0.69 (0.19) | 0.69 (0.18) | 0.69 (0.18) |
|  |  | Gain (%) | - | **-0.3** | **-0.3** | **-0.6** |
| Plant Height | CV2 | Mean | 0.92 (0.01) | 0.92 (0.01) | 0.92 (0.01) | 0.93 (0.01) |
|  |  | Gain (%) | - | **0.0** | **0.1** | **0.2** |
|  | CV1 | Mean | 0.84 (0.04) | 0.84 (0.04) | 0.85 (0.03) | 0.86 (0.03) |
|  |  | Gain (%) | - | **0.2** | **2.0** | **2.1** |
|  | CV00 | Mean | 0.53 (0.20) | 0.54 (0.20) | 0.58 (0.16) | 0.59 (0.15) |
|  |  | Gain (%) | - | **1.3** | **10.2** | **11.9** |
| Grain Yield | CV2 | Mean | 0.75 (0.03) | 0.76 (0.03) | 0.76 (0.03) | 0.76 (0.03) |
|  |  | Gain (%) | - | **0.4** | **0.5** | **0.9** |
|  | CV1 | Mean | 0.69 (0.06) | 0.69 (0.06) | 0.71 (0.04) | 0.72 (0.04) |
|  |  | Gain (%) | - | **0.4** | **4.2** | **4.5** |
|  | CV00 | Mean | 0.52 (0.18) | 0.54 (0.18) | 0.57 (0.15) | 0.59 (0.14) |
|  |  | Gain (%) | - | **3.7** | **9.5** | **13.5** |

We used Model 4 to study the effect of within-population gene diversity (Hs) on grain yield stability across various climatic conditions (Figure 4). The distribution of grain yield G×E best linear unbiased prediction (BLUP_G×E_) across the EVA Network was highly variable across landraces, suggesting that grain yield stability varied among landraces (**Figure 4A**). We then analyzed the relationship between Hs and the standard deviation of BLUP_G×E_ for each landrace predicted by the model M4 (SD_BLUPG×E_). Hs and SD_BLUPG×E_ were negatively linearly correlated (*b* = -0.76, *p* < 0.001, *R*^2^ = 0.191; **Figure 4B**), suggesting that landraces with the highest diversity were more stable across environments. In addition, SD_BLUPG×E_ was also positively correlated with the landrace genetic value (BLUP_G_), suggesting that high performing landraces displayed also higher G×E (**Figure S9**). To test whether these relationships could come from a spurious effect of our sparse design, we first analyzed if we accurately predicted observed SD_G×E_ (**Figure S9**). Predicted SD_BLUPG×E_ was positively correlated with observed G×E (*b* = 1.3, *p* < 0.001, *R*^2^ = 0.05, **Figure S9**) and this correlation increased for landraces evaluated in a high number of environments (b=2.2, r2=0.29 for landraces evaluated in at least 4 environments). This suggests that our model predicted accurately SD_BLUPG×E_, and that, as expected, this accuracy increased with the number of environments and that we certainly underestimated SD_G×E._ (**Figure S9**). In addition, Hs, SD_BLUPG×E_ and observed G×E varied only slightly according to the number of environments in which landraces were evaluated, indicating that the effect of Hs on GY stability did not originate from an unbalanced distribution of landraces with low or and high Hs across our sparse design (**Figure S10**).

**Fig. 4.**
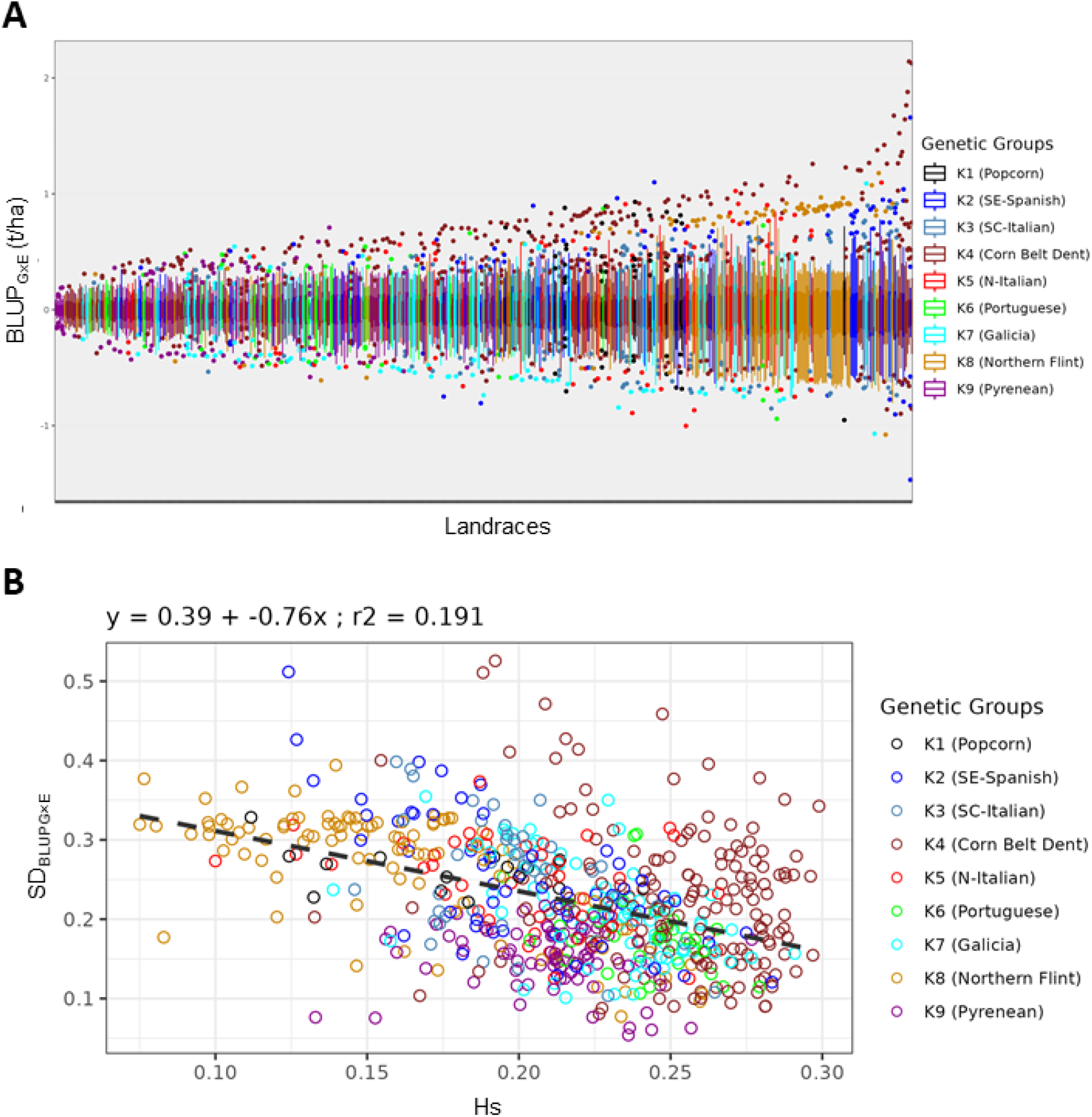
Variation of grain yield stability across field trial network and how it relates with with-in-population gene diversity. **(A)** Variability for each landrace across 22 environments of grain yield genotype-by-environment interaction best linear unbiased predictors (BLUP_G×E_). **(B)** Relationship between within-population genetic diversity (Hs) and grain yield stability assessed using the standard deviation of BLUP_G×E_ (SD_BLUPG×E_). BLUP_G×E_ and SD_BLUPG×E_ were estimated using model 4 that included Hs and GO as covariables.

We then compared 4 different previous models (M1, M2, M3, M4) for their ability to predict GY, PH and MF of landraces across 25 environments using 3 cross-validation schemes corresponding to 3 different prediction scenarios (**Figure S11**): (1) predicting a missing observation (unobserved landrace-environment combination) in a sparse experimental network (CV2), (2) predicting a new landrace (never observed in any trial) in observed environments of this network (CV1), and (3) predicting a new landrace in a new environment by hiding in the calibration set a trial of this network (CV00).

Predictive abilities (PA) obtained for the different traits were highest with the CV2 scenario, followed closely by CV1 (**Table 3, Figure 5**). PA dropped strongly for all traits and models when moving from CV1 to CV00. For instance, PA decreased from 0.89 to 0.69 for MF, from 0.84 to 0.53 for PH and from 0.69 to 0.52 for GY with model 1 (**Table 3**). Adding Hs and GO in the models M2, M3 and M4 increased PA for GY and to a lesser extent PH under CV1 and CV00 scenarios, while it did not increase PA for MF (**Figure 5**). The highest gain in PAs was obtained with M4 under CV00 scenario (new landraces in new environments) with 11.9 and 13.5% of PA gain for PH and GY, respectively compared to M1 (**Table 3, Figure 5**). Including Hs instead of GO in the model led to a higher gain in PA in CV00 scenario for PH (10.2 vs 1.3%) than for GY (9.5 vs 3.7%). For PH and GY, the gain in PA for M4 (Hs + GO) relative to M1 could be obtained by adding the gain in PA of M2 (Hs) and M3 (GO), confirming that Hs and GO contributed independently to trait variation. To test whether our GP models predicted more than the genetic structure, we compared PA obtained under M1 with prediction using a multiple linear regression based on assignments to the nine genetic groups (Table S1, Balconi, Galaretto et al., 2024). M1 outperformed multiple regression based on genetic structure for all traits and all repetitions under CV2, and PA variability was strongly reduced across repetitions (**Figure S12**).

**Fig. 5.**
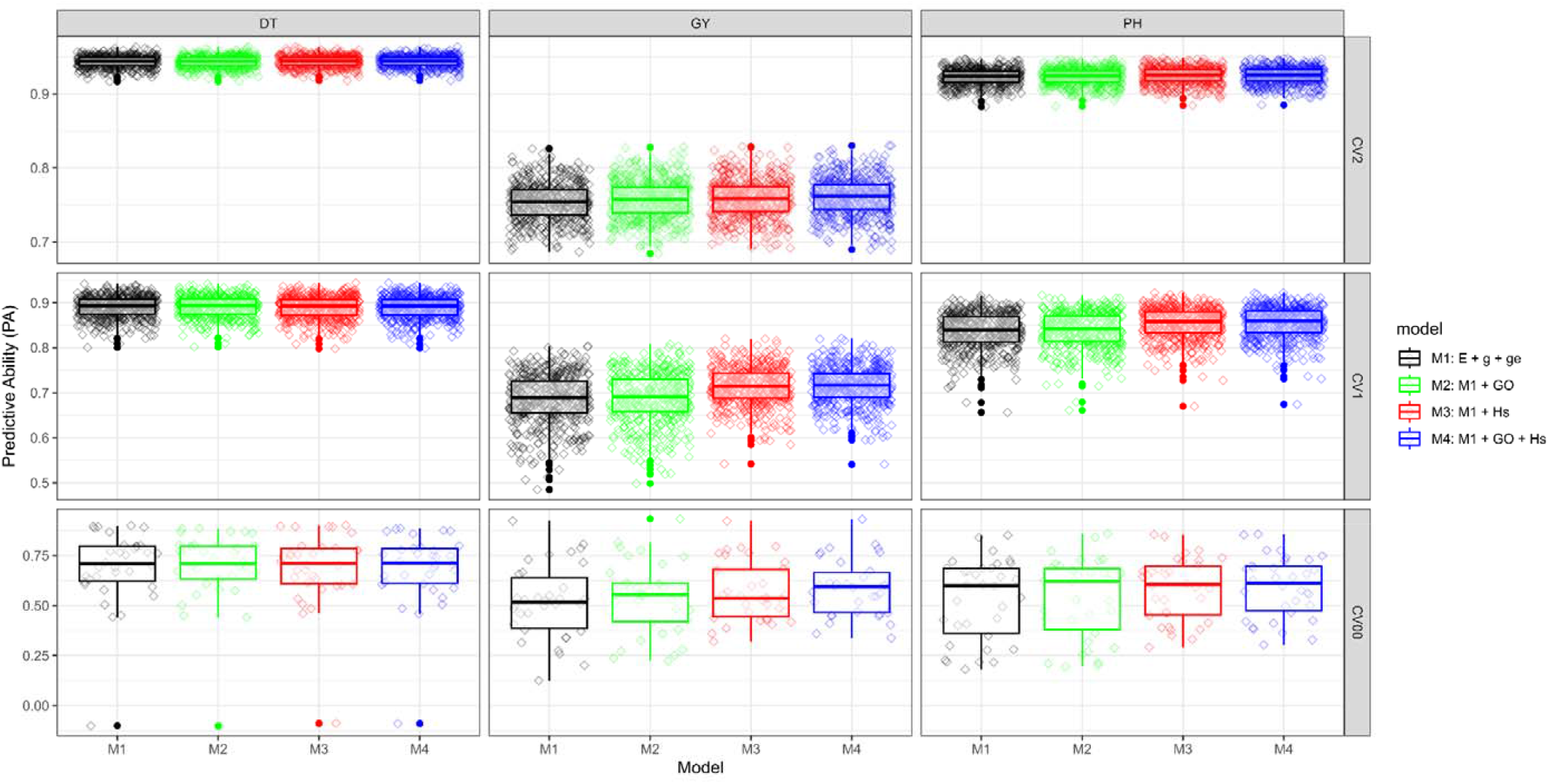
Predictive ability for models and traits. Predictive abilities obtained using different genomic prediction models for male flowering (**MF – left**), grain yield (**GY - middle**) and plant height (**PH – right**) under **CV2 (top)**, **CV1 (middle)** and **CV00 (bottom)** cross validation scenarios. Under **CV2**, 20% of phenotypic observations (genotype-environment combinations) were randomly removed from the whole training set then predicted using 80% remaining training set. Under **CV1**, 20% of the genotypes were randomly selected and their phenotypic observations were removed across the dataset; as a consequence, genotypes to be predicted were not present in the training set. Under **CV00**, one trial is removed at each fold and their genotypes were removed also across other trials; as a consequence, both environment and genotype to be predicted were not present in the training set. Each point represents the correlation between observed and predicted values for one cross-validation fold.

PA distribution showed that PA increase for PH and GY in M3 and M4 was mostly driven by PA increase of the worst predicted environments, leading to a decrease in PA variance (**Figure 5**). In particular, PA augmented in the three environments of Pontevedra (COS_2023_1B, CES_2022_1B, and CES_2023_1B) for GY from 0.13, 0.26 and 0.29 in M1 to 0.51, 0.40 and 0.53 in M4, respectively (**Figure S13**). The same trend was evidenced for one experiment at Delley (DDS_2024_2B3B), where PA increased from 0.20 in M1 to 0.30 in M4. This could indicate that incorporating GO is most valuable for predicting GY in new environments, that are difficult to predict because they differ from the ones included in the training set.

### Integrating genomic offset and genomic prediction enabled the identification of promising landraces adapted to future European climates

We investigated how to use the GF model in combination with the GP model integrating GO and Hs to identify landraces that could be adapted to future climatic scenarios. To do so, we devised a three-step scenario for selecting landraces that are well adapted (low GO), have a high grain yield (high genetic value) and are stable in fluctuating environments (low SD_BLUPG×E_) under a pessimistic (SSP585) long term (2081-2100) future climatic scenario. In a first step, we used GF model for selecting a list of landraces in the EVA panel with low GO (<0.03) under this future scenario conditions for two relevant target regions for maize production in Europe: NE-Romania and SW-France (**Figure 6**).

**Fig. 6.**
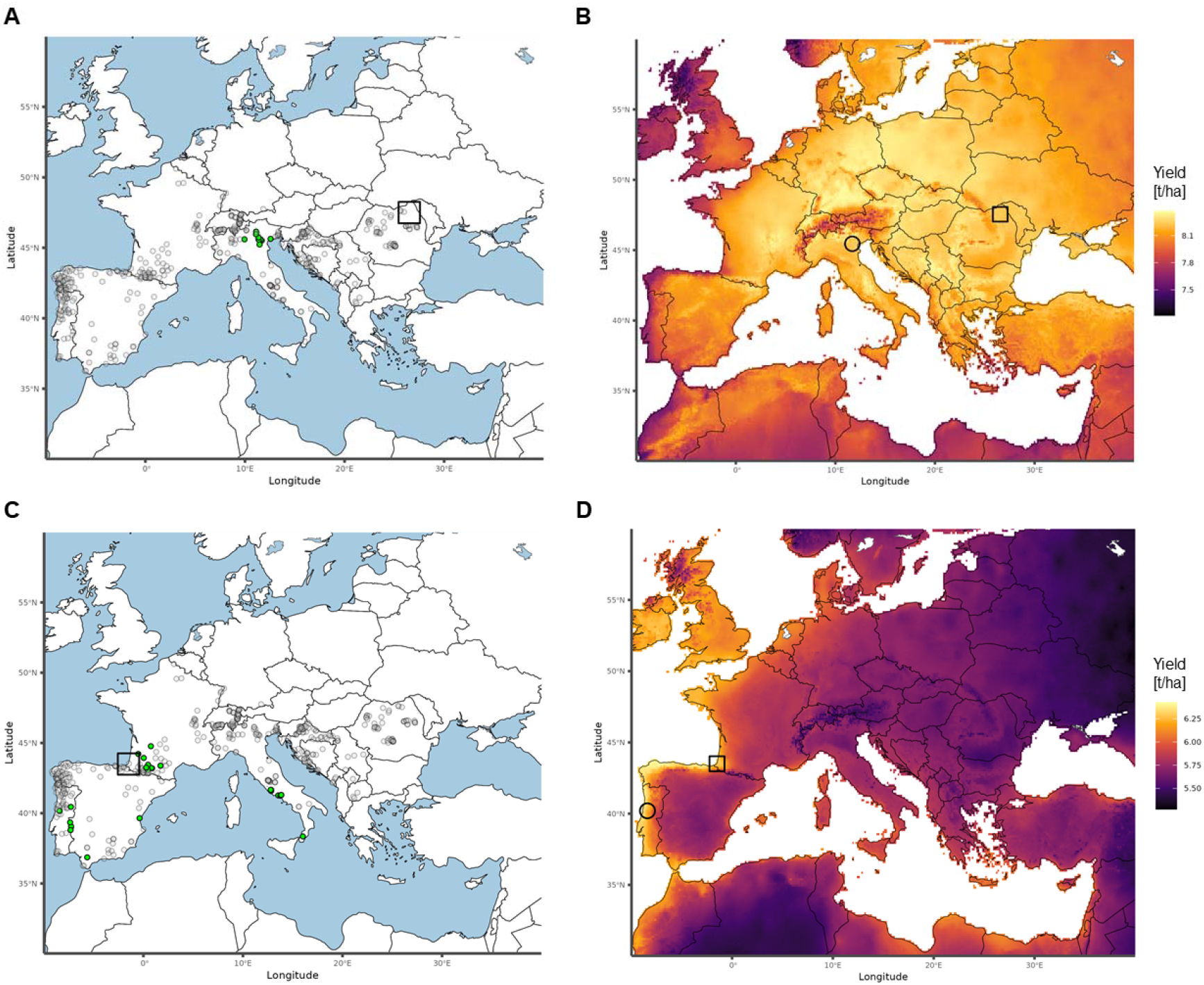
Interesting landraces for future climatic scenario and yield variation due to genomic offset (GO). Identification of landraces with low genomic offset (GO <0.03; collection sites shown in green) for target regions (squares) of NE-Romania **(A)** and SW-France **(C)** under climatic conditions of SSP585 climate scenario for 2081–2100. Yield variation due to GO under this scenario is projected for two landraces selected considering predicted grain yield and stability, (ITA0709 and PRT0537) originating from countries different from the target regions (**B** and **D**, collection sites represented by circles). This example illustrates how combining GP with GO can support the identification of exotic genetic resources that satisfy different criteria.

Across the EVA Panel, we identified 10 and 27 landraces presenting low GO for these two regions, respectively (**Tables S8-S9**). For NE-Romania, local landraces had high GO estimations (**Figure 6A**) while for SW-France, local landraces had low GO in future scenario (**Figure 6C**). This suggests that Romanian landraces will be maladapted under this future scenario and a stronger need for looking for non-local resources to breed for adaptation. In a second step, we used GP model integrating GO and Hs (M4) to predict agronomic performance of these landraces and to assess their yield stability using SD_BLUPG×E_ (**Figure S14, Tables S8-S9**).

For NE-Romania, predicted grain yield of selected landraces varied between 1.9 and 4.2 t/ha, while SD_BLUPG×E_ varied between 0.18 and 0.31 t/ha. None of these landraces were among the top 20% for both yield and stability (low SD_BLUPG×E_) (**Table S8**). For SW-France, predicted grain yield of selected landraces varied between 0.4 and 2.6 t/ha, while SD_BLUPG×E_ varied between 0.08 and 0.40 t/ha. Two landraces were among the top 20% for both yield and stability (**Table S9**). Following these two criteria (the highest performing and most stable), we identified PRT0537 and ITA0709 as promising landraces for NE-Romania and SW-France, respectively (**Figure S14**).

We then illustrated yield variation of these two landraces across Europe in this future scenario. The Portuguese landraces PRT0537, displayed the best yield along the western coast of Portugal and Spain, in addition to SW France (target region) (Figure 6B). The Italian landrace ITA0709, displayed a good yield performance across all Europe except the Western Atlantic coast and the mountain regions in these future scenarios (Figure 6D). This landrace showed a low loss of performance for most part of the SE-European agricultural region, but a higher predicted yield loss due to increased GO in western France and most of Spain and Portugal.

We then investigated whether some landraces will be favored by climate change or not in regions across Europe with genomic prediction model M4. We estimated yield variation due to GO for four landraces representing different genetic groups and how it will evolve (yield gain/loss) in SSP585 at 2081-2100 as compared to the past climate. Whatever the landrace, we observed a higher yield loss in the region of origin of the landrace (or in regions with similar environment) which is expected since GO could only decrease in this region. Interestingly, yield change strongly depends on the landraces. For instance, the model predicted that the yield of Southern Spanish landraces (such as ESP0148) will strongly increase in the North of Europe above latitude 47° (>1t/ha) where they were not originally adapted (**Figure S15**). Similarly, the French Pyrenean landrace FRA0491 will lose yield in Western France but will gain yield in Germany and Eastern Europe. In contrast, no yield gain was observed for the Bosnian Landrace BIH0063 or the Northern Eastern French landrace FRA0296 suggesting that these landraces will be no longer adapted to any European region in the future. Taken together, our results showed that climate change may differently impact landraces, according to their region of origin and whether the environment corresponding to the landrace collection site will expand or reduce in the future.

## Discussion

### HPG-based genomic prediction is an efficient and cost-effective approach to characterize genebank collections of landraces

Exploiting genetic resources and adaptive alleles is essential for addressing future agricultural challenges (Godfray et al., 2010; Hoisington et al., 1999). Given the vast number of accessions preserved *ex-situ* in gene banks, efficient tools are needed to identify the most promising ones for pre-breeding programs (Böhm et al., 2017; Bohra et al., 2022; Gorjanc et al., 2016; Mayer et al., 2020; *FAO*, 2025). In our study, we investigated the potential of combining high throughput DNA-pool genotyping (HPG), genomic prediction (GP), genomic offset (GO) and within-population gene diversity (Hs) to predict performances of maize landraces for agro-morphological traits and their adaptation to future climatic scenarios. HPG enables to significantly reduce genotyping costs, becoming a feasible strategy for massive characterization of genebank collections, especially for heterogeneous population (Arca et al., 2021; Mascher et al., 2019).

Our GP models trained with a sparse evaluation network accurately predicted landrace agro-morphological traits in observed environments, in which they never were evaluated (prediction scenarios “CV2” and “CV1”). This corresponds to a typical genebank use case in which we would like to predict the phenotype of an unobserved landrace across various environments based only on their genotype and a training set of landraces that were both genotyped and phenotyped (El Hanafi et al., 2023; Keep et al., 2020; Mascher et al., 2019). Regarding traits, MF showed the highest predictive ability, followed by PH and GY, whatever the prediction scenario. These results are consistent with higher heritabilities and lower genotype by environment (GxE) effects for MF than for PH and GY (Hallauer et al., 2010). Integrating GO and Hs into the GP models was most useful for improving GP accuracy for the most challenging prediction scenario (predict a new accession in a new environment) and the most challenging trait (low heritability and high G×E) (**Table 1-3**).

Regardless of the model used, GP was robust and efficient in predicting landrace trait across environments under a sparse design (CV2), yielding high PA (0.75-0.94 depending on model and traits, Table 4). This result is in agreement with PA obtained through HPG-based GP in forage species (Cericola et al., 2018; Guo et al., 2018; Keep et al., 2021; Pégard et al., 2023). Including GO and / or Hs neither improved nor reduced PA in this scenario, suggesting that Hs and GO effects were already captured by genetic and G×E effects in this sparse design scenario. Under CV2 scenario, the prediction of missing landrace-environment phenotypic values is enabled by correlated information from other environments and related landraces (Burgueño et al., 2012a).

Combining HPG-based GP with a sparse evaluation networks for evaluating traits such as the ECPGR EVA Maize Network (Balconi, Galaretto et al., 2024) is therefore an efficient and cost-effective strategy to characterize agro-morphological traits in various environments (**Table 3**). In addition, the phenotyping cost could be reduced without affecting PA by optimizing training set of landraces and the field trial network for capturing G×E interaction. Previous works suggest that it could be possible to optimize the training set composition of landraces based on their genotyping without affecting PA but this approach remain to evaluate on landraces (Pégard et al., 2023; Rincent et al., 2012). Our result also paves the way for reusing thousands of historical and patrimonial phenotyping experiments and field observations of maize landraces made by genebanks and research teams to train GP models and therefore valorize this tremendous historical work. However, it required to genotype these landraces and to digitalize this patrimonial field experiment.

### Within-population gene diversity contributes positively to agronomic performance and stability

Within-population gene diversity (Hs) had a strong positive effect on GY and PH but not on MF. Hs captured 8% and 20% of PH and GY genotypic variation, respectively (Table 2). This positive correlation of Hs with GY and PH is consistent with previous studies based on RFLP markers on maize landraces (Rebourg et al., 2001). Although this effect is widely recognized in maize, it is rarely included in GP models. Accounting for inbreeding depression in GP of maize hybrids at individual level improved PA for height and yield (Roth et al., 2022). Our study showed that this result obtained at individual plant level could be extended at the population level to improve PA of landraces cultivated *per se*. For PH and GY, including Hs increased PA when predicting unknown landraces in known (CV1) and unknown (CV00) environments. The Hs effect was independent from GO effect, suggesting that Hs is mostly related to the potential intrinsic average performance of landraces while GO is mostly related to G×E. In addition, our results suggested strongly that landraces with higher Hs had more stable GY across the EVA field trial network, suggesting that within-population gene diversity partially determines the capacity of landraces to buffer fluctuating environmental conditions. This buffering environmental effect of within-population gene diversity was observed in several cultivated species and studies when comparing the yield of cultivar mixtures to monoculture, since the yield of cultivar mixtures outperformed monocultures, with a stronger effect in environments with biotic and abiotic stresses and in years with extreme weather (Reis and Drinkwater, 2017). This buffering effect could be due to the fact that landraces with larger population genetic diversity have more chance plants that are adapted to a given environment compensating the detrimental effect of plants that are not adapted but also that this heterogeneity limits disease propagation.

The Hs effect certainly originates from the well documented strong effects of inbreeding depression on these traits in maize (Hallauer et al., 2010) driven by exposing deleterious alleles or by decreasing the number of best-fitted heterozygotes (Crow & Kimura, 1970). In a random mating outcrossing population, Hs corresponds indeed to the expected mean heterozygosity of plants and therefore relates directly to their level of inbreeding. Low within-population gene diversity for some landraces maintained *ex-situ* could originate from several factors that reduce the effective size of the populations, thus creating a bottleneck. First, some landraces may have undergone loss of diversity before their collecting because they have been cultivated in small fields or gardens in which the number of plants was low. Second, they may have been collected initially with a number of ears that was too low to well represent landrace diversity. Third, not enough plants may have been used for regenerating landraces seed-stock in genebanks.

Whatever its origins, low within-population gene diversity has a very strong effect on decreasing agronomic performances, notably for yield and plant height. Farmers must account for this inbreeding depression when cultivating and multiplying landraces, particularly when the field area is small. Inbreeding could be avoided by reintroducing genetic diversity by crossing with landraces that have similar agro-morphological characteristics coming from genebanks or other local farmers through seed exchange. The admixture between landraces from different genetic groups should mitigate inbreeding depression effects, improve performance at the population level and enhance adaptive potential required to cope with a changing climate (Aguirre-Liguori et al., 2019).

### Genomic offset promises for maize landraces: some things to win, few things to lose, many to explore

Including GO increased PA of GY and PH only when both genotype and environment were unknown (CV00), capturing a part of G×E for these two traits (Table 2, Figure 4). As GO effect was negative, these results confirm the hypothesis that maize landraces are locally adapted and that maladaptation increases as climatic conditions diverge from those of the collection site. Our study likely underestimates the effect of GO on traits since only 10% of the landraces have been evaluated in field experiments with contrasted GOs (Figure S4). We observed that GO had no impact on PA of MF, consistent with lower magnitude of G×E for this trait (Hudson et al., 2022; Rogers et al., 2021). Contrary to our study, (Li et al., 2025) reported that climatic conditions of landrace collection site have limited importance when predicting landrace value for breeding. These contrasting findings are likely due to the fact that our study addresses *per-se* performance of landraces, while signals of adaptation may be blurred by testers in gamete capture experimental designs (Li et al., 2025). Secondly, we used DNA-pool genotyping approach which better captures within-population gene diversity while Li et al., 2025 used a single plant genotyping that could blur the relationship between allelic frequency and environmental gradient. In addition, GO signal of adaptation may be reinforced in our study by the use of gradient forest (GF) that ranked, selected, weighted climatic variables according to their effect on allelic frequency (Beck et al., 2025).

Nevertheless, the ability of the GO to capture G×E could be improved further by tackling several limiting factors from our study. First, our sparse testing design does not allow to optimally capture G×E effects for all landraces, since only a small set of landraces have been evaluated in contrasted environments. Thus, more complete designs should improve G×E quantification. Secondly, our GO analysis relied on full-year climatic variables, which coarsely represent the actual growing conditions at field experiments and collection sites. Replacing satellite-derived climatic data by local climatic records, and accounting for management practices (e.g., planting date, irrigation) and plant phenology will certainly improve GO measurement. Moreover, including additional environmental descriptors—such as solar radiation, evapotranspiration, or soil properties could further improve environmental characterization and therefore the measurement of GO. Ideally, experimental designs aiming to establish the relationship between GO and phenotypic traits should target environments that represent contrasting GO conditions (Lotterhos, 2024). Consequently, landraces would likely be exposed to both high and low GO conditions, improving the assessment of GO effects. Accordingly, using a subset of landraces with contrasted GO exposure in the field network increased strongly GO effect on GY and PH (**Figures S7**). Complete experimental designs would further enable the study of genotype-by-GO interactions, as we observed that landraces may differ in their phenotypic responses to GO (data not shown).

Investigating how GO sensitivity relates to landrace history could also provide additional insight for genetic resources. Lack of sensitivity to GO could be related to a weak natural selection pressure due to the variability of climatic conditions at the collection site (Keep et al., 2021) or specific management practices like irrigation that mitigate climatic effects. In addition, *ex situ* multiplication of landraces by genebanks could have led to adaptation to climatic conditions that differ from those at the original collection site. Maize landraces are also subject to human selection for specific purposes, such as earliness illustrated by short-cycle Italian landraces from central-southern regions that are cultivated after wheat or grain quality, as reflected in the marked differences between landraces used for human consumption (e.g., polenta, bread) and those intended for animal feed (birds, monogastrics). (Balconi et al., 2024; Gouesnard et al., 1997; Redaelli et al., 2024). Prior knowledge related to the original use and field management of the maize landraces could help to anticipate the relevance of GO as an indicator of adaptation or its impact in GP model accuracy for certain accessions.

### Identify pre-adapted genetic resources for future climatic conditions by integrating genomic offset and genomic prediction

We also illustrated how GO predictions can be used to identify landraces with potential adaptation to future climatic scenarios. Our model predicted strong negative effects in specific regions of Europe that are consistent with results from different studies using crop-climate models (Heino et al., 2023; Jägermeyr et al., 2021; Knox et al., 2016). The utilization of GO to identify interesting genetic resources has been deployed in similar approaches like “optimal migration” for pearl Millet (Rhoné et al., 2020) or “reverse offset” for forest tree (Gougherty et al., 2021). GO predictions for a target region or climate scenario can provide a reduced list of promising genetic resources but they do not predict traits in these new target environments. A related strategy, the Focused Identification of Germplasm Strategy (FIGS), has successfully identified valuable sources of stress-tolerance variation (Stenberg & Ortiz, 2021) but similarly to GO, this approach does not predict traits of interest in new environments.

Our approach overcome the issue of two previous mentioned strategies by using a GP model integrating GO and therefore assigning a phenotypic scale to the GO change, allowing to provide the phenotypic landscape due to GO. Scaling GO for different traits is a major challenge in eco-genetics approaches that remains to be addressed (Fitzpatrick et al., 2025). Nevertheless, GO effect remains to be validated using adequate field experiments as it may be subject of different limitations (see above). Combining G×E in genomic prediction model with GO is an encouraging way to address this issue.

Our work opens several perspectives for ecological genomics as genomic forecasting of mal-adaptation through GO could be translated into phenotypic change through our prediction model. This prediction model could be improved since GO captured only a part of G×E interactions or environment-specific effects due to limits of our sparse design for evaluating GO relationship with traits for each landrace (see below). Since GO relationship with traits could be different between landraces or genetic groups (data not shown), including a specific landrace or genetic group GO effect rather than an average effect in our model could allow to explore more detailed responses to GO. This would require a complete design rather than a sparse design. Integrating an environmental kinship in our model could also capture more complex G×E interactions that are not captured by GO. Another limitation of our approach relates to (as any GO model) to the need of “geolocated” accessions in order to model and predict (mal)adaptation due to climatic conditions. Genomic prediction of collection site climatic variables (environment of origin) could be a solution, but its contribution to prediction accuracy remains to be explored (Li et al., 2025).

To conclude, our study is the first to apply GP using the HPG approach to characterize the *per se* performance of maize landraces across multiple environments. HPG-GP approach paves the way for low cost characterization of landraces, that are conserved either *ex situ* in genebank or *in situ* by farmers, for various complex traits as phenology, performance, quality traits, tolerance to abiotic stress. This is also the first study to show that GO and Hs can provide valuable, complementary information for predicting maize landrace performance opening an avenue for their integration in GP model for characterizing heterogeneous genetic resources. This integrative approach combining GO with GP could be very useful for pre-breeding activities by identifying a subset of landraces suited for cultivation in specific environments (present or future) based on both an adaptation criterion (GO) and an agronomic performance criterion (GP-GO model). GO predictions highlighted that landraces coming from other geographic area may be pre-adapted to future conditions in regions highly disturbed by climate change, supporting the need of international collaboration to adapt crops to a changing climate. Integrating GO and Hs into the GP models was most useful for improving GP accuracy for the most challenging prediction scenario (predict a new accession in a new environment) and the most challenging trait (low heritability and high G×E). In the end, our GP-GO model also predicted stability of landraces in fluctuating environments which is relevant considering that climatic change will increase climatic instability in the future (Heino et al., 2023).

## Material and Methods

### Plant material

Accessions from a panel of 626 European maize landraces (hereafter called “EVA Panel”) from nine European genebanks, previously assembled and described in Balconi, Galaretto et al. (2024). Landrace selection were made by each genebank primarily to represent national diversity in geographical origin and variability in morphological and agronomic traits, including tolerance to biotic and abiotic stresses. Based on DNA pool genotyping, Balconi, Galaretto et al. (2024) identified nine main genetic groups using Admixture (Alexander et al., 2009): Popcorn (K1), SE-Spanish (K2), SC-Italian (K3), Northern Flint (K4, NF), N-Italian (K5), Portuguese (K6), Galician (K7), Corn Belt Dent (K8, CBD) and Pyrenean (K9) (**Table S1**). Note that the interpretation of K8 as the “Corn Belt Dent” genetic group in Balconi, Galaretto et al., (2024) is based on the quantitative assignment of European landraces to the American “Corn Belt Dent” group using the structuration analysis of 156 landraces from America and Europe (Arca et al., 2023).

We retrieved geolocation for 483 landraces based on their collection sites described in passport data from EURISCO Catalog (Kotni et al., 2023). Using these coordinates, we retrieved climatic variables from climatic databases (see section “climatic characterization” in Methods). Finally, a subset of 470 landraces with no missing climatic data was used to model adaptation using gradient forest (GF) (Ellis et al., 2012; see section “genomic offset” in Methods).

### Genotyping and diversity analysis

The landraces from the EVA panel were genotyped with the 50K Illumina Infinium HD array (Ganal et al., 2011) and allelic frequency was obtained for 35,878 SNPs, following a DNA pool approach proposed by Arca et al. (2021) and analyzed by Balconi et al. (2024) for structure analysis. Markers without missing data and minor allelic frequency higher than 0.05 were retained. Out of the 31,950 SNPs, after filtering, a subset of 23,412 SNPs with low ascertainment bias was used to estimate gene diversity within each landrace (within-population gene diversity, Hs) and genetic relatedness (kinship) between landraces.

We retrieved the Hs estimation made by Balconi, Galaretto et al. (2024) following Nei (1973). We estimated the Kinship matrix (*K*) using allelic frequencies according to Cericola et al. (2018), as:

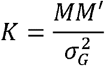

where *M* is the centered allelic frequency matrix, *M’* is transposed matrix of *M* and 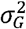 is the sum of expected variances across SNPs, defined by Ashraf et al. (2014), as:

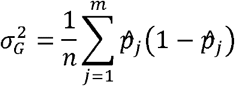

where 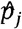 is the allelic frequency of *j*^*th*^ SNP at the whole panel level, *m* is the total number of SNPs and *n* is the ploidy number which is equal to 2 for maize.

### Analysis of agronomic traits evaluated across EVA field trial Network

The European Evaluation Network (EVA Network) was established to evaluate a large number of maize landraces in field experiments under a sparse testing design (not all landraces were evaluated in all experiments). Targeted traits involved phenology, agro-morphological characters and tolerance to biotic and abiotic stress. A total of 64 experiments were carried out between 2021 and 2024 across Europe where *per se* value of 588 landraces were evaluated along 5 commercial hybrid checks. Experiments were conducted by genebanks, research institutes and private partners under the framework of the EVA Network led by the European Cooperative Program for Plant Genetic Resources (ECPGR - https://www.ecpgr.org/eva/eva-networks/maize).

We used a subset of 397 landraces evaluated for grain yield, complete collection site climatic information and within-population gene diversity above 0.05 (see next section in Methods) to evaluate the ability of genomic prediction (GP) to predict landraces traits across 25 different environments (**Table S1**). All landraces had been evaluated for plant height (PH), grain yield (GY) and days from sowing to male flowering. We estimated the thermal time for male flowering (MF) as growing degree units (GDU) accumulated from sowing to when 50% of the plants have shed pollen, using Tb = 6°C as in Rebourg et al., (2001). Plant height (PH) was recorded as the average of measurements taken from 10 competitive plants per plot. For each plant, PH was measured in centimeters as the distance from ground level to the base of the tassel.

We determined GY as the weight of shelled grain from all harvested ears per plot, adjusted to 10% grain moisture and expressed in t/ha. The 25 environments involved 6 locations: Belgrade (Serbia), Coimbra (Portugal), Avenches (Switzerland), Pontevedra (Spain), Suceava (Romania) and Zagreb (Croatia), four years (2021, 2022, 2023, and 2024), and different evaluation conditions: standard (*n* = 19), early sowing (*n* = 2), inoculation with the fungus *Fusarium verticilloides* (*n* = 2), and artificial infestation with the pest *Sesamia. nonagrioides* (*n* = 2) (Table S6). The number of entries and experimental design varied across the network (**Table S5 and Table S6**). Landraces were evaluated on average in 2.8 environments, with five landraces evaluated in a maximum of 12 environments (Table S5). The number of landraces per environment varied from 17 to 165, with an average of 45 (**Table S6)**.

We corrected field plot values for spatial and design effects in each environment, as by Roth et al. (2022), using the mixed model of the form:

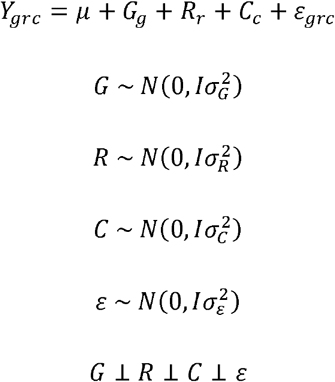

where *Y*_*grc*_ is the plot value of genotype *g* (landrace or check) measured at row *r* and column *c* and 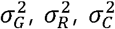 and 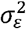 are genotypic, row, column and error variances components, respectively. All effects were considered random and independent (⊥). Heritability for each trait in each environment was estimated as 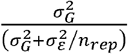, with the number of replications (*n*_*rep*_) varying between 1 and 3 (**Table S5**). We estimated corrected field values 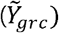 by removing the spatial and design effects from the plot values. Then, we computed adjusted means within environment 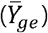 as the mean of the corrected field values. Finally, we estimated least square means (*lsmeans*) across environments for each landrace 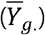 with the model:

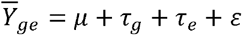

where *τ* _*g*_ and *τ* _*e*_ are the fixed effects of genotype *g* and environment *e*, ε is the random error. The *lsmeans* for each genotype *g* was estimated as 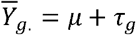

### Climatic characterization

To characterize climatic conditions in which landraces were collected, we used original collection site coordinates to retrieve past climatic information from WorldClim2.1 (Fick & Hijmans, 2017). WorldClim2.1 provides 55 climatic variables, including monthly averages of maximum temperature (*tmax*), minimum temperature (*tmin*), precipitation accumulation (*prec*) and 19 bioclimatic variables (*bio1* to *bio19*) for the period 1970-2000 (see **Table S2** for definitions). We used R packages *raster* (Hijmans, 2010) and *terra* (Hijmans, 2020) to extract climatic and bioclimatic variables for the 483 collection site coordinates available. After removing sites with missing values, a total of 470 landrace collection sites (**Table S1**) and 55 climatic variables were retained for further analyses. Summary statistics for each climatic variable are provided in **Table S2**.

In order to characterize climatic conditions at environments (field experiments) of the EVA Network we used NASA Power (Sparks, 2018) to retrieve daily information on temperature and precipitation. To enable comparison of these climatic conditions with those at landrace collection sites, we compiled temperature and precipitation data on a monthly basis and used *biovars* function of *dismo* R package (Hijmans et al., 2010) to obtain the same bioclimatic variables than WorldClim2.1. Summary statistics for climatic data of EVA Network are also provided in Table S2.

### Genomic offset estimation using gradient forest

Genomic offset (GO) is a metric of maladaptation based on the prediction of disruption of (assumed) optimal gene-environment relationship (Fitzpatrick & Keller, 2015). In order to estimate GO between the climatic conditions experienced at landrace collection sites and the environments included in the EVA Network, we first modeled the change in allelic frequencies across climatic gradients using the R package *gradientForest* (Ellis et al., 2012). Gradient forest (GF) is a Random Forest-based algorithm that was designed to model the response of biodiversity to environmental gradients. Fitzpatrick & Keller (2015) proposed using genomic data instead of species count to model the response of a set of loci to environmental gradients. This GF model is used to predict optimal allelic frequencies in a given environment, based on environmental variables for current, past or future climates. GO then measures maladaptation as the shift between the allelic frequencies of each population and those predicted by the model for each environment. To select loci involved in adaptation, we first adjusted a GF model including all 55 climatic variables and 35,878 SNPs, using R functions from “geneticOffsetR” (Fitzpatrick et al., 2021), which allowed to fit GF models for a large number of SNPs and recover only GF model goodness-of-fit (R^2^) for each marker. Then, empirical *p-values* were calculated by determining the rank of the R^2^ value for each SNP, within a distribution of intergenic SNPs (Fitzpatrick et al., 2021). For this, we used the set of 2,500 unlinked SNPs previously selected in Arca et al. (2023). 243 outlier adaptive *loci* were retained (p-value < 0.01) and then used in a second GF model to select the most important among the 55 climatic variables WorldClim2.1. We used GF variable importance ranking based on their efficiency for predicting allelic frequency variations, to select the 15 top variables (importance >0.6, **Table S3**). In the end, we fitted a third GF model using these 15 selected climatic variables and these 243 outlier SNPs. This model served two main purposes: first, to analyze the spatial variation in predicted genetic compositions and second, to predict landrace GO for climatic conditions of each EVA Network field experiment.

To achieve this first objective, we used cumulative importance curves derived from GF. These curves represent transformations of the environmental data that are shaped by genomic patterns. They illustrate both the rate and magnitude of allele frequency shifts along each environmental gradient, including any threshold responses. By rescaling the environmental data into common units of genetic turnover, these functions provide a more accurate representation of genetic composition patterns than raw, untransformed environmental data. To draw a geographical map of the impact of environmental gradient on genomic change (turnover map), we first performed a PCA analysis based on GF-transformed climatic variables from each WorlClim2.1 grid across Europe (2.5 arc minutes resolution ∼ 4.5km x 4.5 km at the Equator). We created a multivariate color scale, by assigning Red, Green and Blue to the first three most important PCA axes and applied the resulting color to each grid of the map. The turnover map colors represent patterns of genetic composition, where change in color (turnover region) represent a shift in adaptive allelic frequencies. To investigate the climatic variability of the landrace collection sites and compare them with the environments explored in the EVA network, we projected each collection site and experiment on the different PCA axis, using GF-transformed climatic variables. To facilitate visualization, we removed admixed landraces (assignment to genetic groups <0.6).

Finally, we used the resulting multidimensional rescaled environmental spaces to quantify genetic differences, measured as genomic offset (GO). This was calculated using the Euclidean distance (using the *dist* function of R package *stat* (R Core Team, 2024)) between the population’s original environment and the EVA Network field experiments, following the approach described by (Fitzpatrick et al., 2021). High values of GO indicate large differences between optimal and landrace genetic composition, which is expected to lead to a loss of fitness.

### Variance decomposition of genomic offset and within-population gene diversity effects on traits

To investigate the relationship between GO values across the EVA Network environments and landrace traits, we fitted linear regressions between GO and the within environment adjusted means 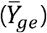 for MF, PH and GY. Similarly, we investigated the relationship between Hs and *lsmeans* 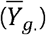 for each landrace across the EVA Network

To quantify the amount of variation explained by GO and Hs, we <u>performed</u> variance decomposition by adjusting different models to the complete data set of corrected field values 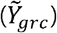. A total of 4 models were adjusted using the *sommer* R package (Covarrubias-Pazaran, 2016) as follows:

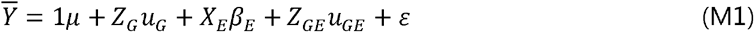

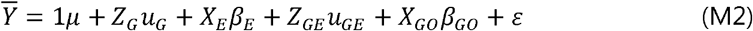

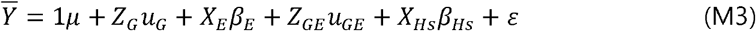

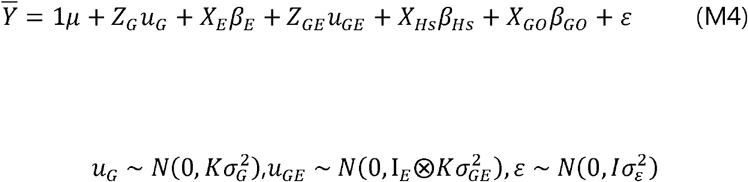

where 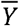 is the vector of phenotypes, 1 is a vector of ones, *μ* is the intercept, *X*_*E*_ is the incidence matrix for the environment fixed effects, *β*_*E*_ is the vector of environment fixed effects, *u*_*G*_ is the vector of main random genetic effects for landraces, *u*_*GE*_ is the vector of random effects for genotype-environment interactions, *ε* is the vector of random errors, *Z*_g_ is the genotypic incidence matrix of size, *K* in the kinship matrix, *Z*_*GE*_ is the genotype-environment interaction incidence matrix, *I*_*E*_ is the identity matrix, *X*_*GO*_ is the vector of GO estimates for the corresponding landrace and environment, *β*_*GO*_ is the GO fixed effect, *X*_*Hs*_ is the vector of within-population gene diversity, *β*_*Hs*_ is the within-population gene diversity fixed effect, ⊗ is the Kronecker product and 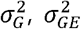 and 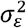 are the unknown genotypic, genotype-environment interaction and error variances, respectively.

### Genomic Prediction

We evaluated the performance of the 4 models for genomic prediction under three different prediction scenarios represented by previously used cross validation (CV) schemes (Burgueño et al., 2012b; Jarquín et al., 2017)(Figure S11):

- **CV2** (prediction under sparse testing): five-fold cross validation where genotype-environment combinations were randomly divided in 5 and at each round four folds (80%) were used for training and the remaining fold (20%) is used for validation
- **CV1** (prediction of new genotypes): five-fold cross validation where genotypes were randomly divided in 5 and, at each round, four folds (80%) were used for training and the remaining fold (20%) is used for validation
- **CV00** (prediction of new genotypes in new environments): at each fold training sets were built using information of all environments, except one that was used for validation. In addition, the data for the genotypes present in the validation set were removed across environment in the training set. Therefore, neither the environment nor the genotypes to be predicted were observed in the training set.

For CV2 and CV1, we repeated random sampling for the five folds 100 times. For CV00, we divided the three biggest environments in more than one-fold to avoid bias related to environment size. As a result, CV00 included 34 different folds of similar size.

We computed predictive ability (PA) for each fold as the correlation between observed and predicted values, as well as mean and standard deviation of PA. Finally, the gain in PA (%) was computed as the relative PA using M1 as the reference.

### Genomic offset application to future climatic scenarios

To investigate the prediction of adaptation or maladaptation of landraces to future climates, we used the GF model to perform GO predictions for future climatic scenarios. We retrieved downscaled CMIP6 2.5-min resolution climatic predictions for two Shared Socio-economic Pathways (SSP126 and SSP585) and two time periods (2041-2060 and 2081-2100) from WorldClim2.1 (https://www.worldclim.org/data/cmip6/cmip6climate.html). We computed GO individually for each of the 13 Global Climatic Models (GCMs, **Table S4**) to then obtain the mean landrace GO for each future scenario (SSP-time period combination). We used mean landrace GO for each scenario to visualize GO landscape and evolution by mapping GO to each 2.5-min grid in the map, using the *ggplot2* R package (Wickham et al., 2007). To provide representative examples, we selected 4 landraces from different geographic regions and genetic groups and represented GO at two periods in the future (2041-2060 and 2081-2100) using these different scenarios of global warming (SSP126 and SSP585).

We then used the GO values to screen among the landraces for putatively best adapted ones (GO <0.03). We focused on two relevant maize production regions of Europe: NE-Romania and SW-France. Among landraces with low GO, we selected landraces from different regions as diversity source, considering performance and stability for grain yield derived from GP M4 (GO+Hs). We assessed yield using the genotypic best linear unbiased predictor (BLUP_G_), Hs effect and GO effect, while yield stability was assessed using the standard deviation of G×E BLUPs (SD_BLUPGxE_). Before incorporating SD_BLUPGxE_ as a stability indicator, we studied the relationship of SD_BLUPGxE_ with Hs, BLUP_G_, and observed standard deviation G×E (SD_GxE_), to discard possible spurious relationships. We computed observed SD_G×E_ as the standard deviation of observed GxE effect in each environment: 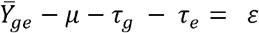 Then, we selected one landrace for each target region considering predictions of yield and stability, and used the GO effect retrieved from M4 to generate yield variation map across Europe linked to GO.

Finally, we assessed the consequences of climate change on landrace performance under the most pessimistic scenario (SSP585 – 2081-2100). We computed the differences between past and future GO across Europe and we used the GO effect to transform GO difference into yield losses/gains.

## Author Contributions

Conceptualization: AOG, SDN, AC, BG, LM

Methodology: AOG, SDN, LM, MP, AC, BG

Investigation: DMa, CV, CBau, RM, AB, PR, CBal, PMM, HS, AMB, DMu, BSC, AS, VA, SG

Formal analysis: AOG, SDN, AC, LM, MPVisualization: AOG, MP, SDN, AC, LM, MP

Supervision: SDN, AC, LM, BG

Writing original draft: AOG, SDN

Writing review & editing: AOG, SDN, AC, LM, RM, CBal, AB, PR, HS, VA, SG

Project administration: SG Funding acquisition: SG, SDN

## Data Availability

Information on accessions and environmental data are all available in supplementary materials. All data required for statistical analysis (Genetic structuration, Genotyping, Phenotyping and Environmental data) are also available in dataverse https://entrepot.recherche.data.gouv.fr/ with following name “EVA Maize genotyping and phenotypic dataset” (https://doi.org/10.57745/TNAURQ) and with a private link for peer-reviewing (https://entrepot.recherche.data.gouv.fr/privateurl.xhtml?token=5ad9e67e-1772-4980-ac4e-28475cd4f4f7). Raw phenotyping data are available on request and will be made available in the EURISCO catalogue (www.ecpgr.org/eurisco). Seeds from each accession can be supplied on request by the different European genebanks (holding institute) listed in the accession list (Table S1), in accordance with the FAO Standard Material Transfer Agreement.

## Declaration

All authors declare that they have no competing of interest. Dr. Alexandre Strigens is employed by the company DSP—Delley Semences et Plantes SA, Switzerland. All authors declare that the research was conducted in the absence of any commercial or financial relationships that could be construed as a potential conflict of interest.

## Acknowledgments

We acknowledge Tristan Mary-Huard for his help with statistical model notation. We are grateful to the INRAE MIGALE bioinformatics facility (MIGALE, INRAE, 2020. Migale bioinformatics Facility, doi: 10.15454/1.5572390655343293E12) for providing computing and storage resources.

We acknowledge the European Cooperative Programme for Plant Genetic Resources (ECPGR, www.ecpgr.org) and all EVA partners for making the European Evaluation network possible and for encouraging the use of genetic resources. We are thankful to the European genebanks within the EVA Maize network and their curators for supplying passport data and landrace’s seeds for both genotyping and field evaluations: University of Zagreb, Faculty of Agriculture, Croatia; INRAE - Centre de Montpellier (CRB-GAMéT), France; Centro di Ricerca Cerealicoltura e Colture Industriali (CREA-CI), Sede di Bergamo, Italy; Banco Português de Germoplasma Vegetal (BPGV-INIAV), Portugal; Escola Superior Agrária de Coimbra, Portugal; Suceava Genebank, Romania; Maize Research Institute Zemun Polje, Serbia; Misión Biológica de Galicia (MBG-CSIC), Spain; Agroscope, Switzerland. We also thank all public and private partners within the EVA Maize network for their contributions to evaluations and fruitful discussions.

We acknowledge the World Climate Research Programme, which, through its Working Group on Coupled Modelling, coordinated and promoted CMIP6. We thank the climate modeling groups for producing and making available their model output, the Earth System Grid Federation (ESGF) for archiving the data and providing access, and the multiple funding agencies who support CMIP6 and ESGF.

## References

Aguirre-Liguori, J. A., Ramırez-Barahona, S., Tiffin, P., & Eguiarte, L. E. (2019). Climate change is predicted to disrupt patterns of local adaptation in wild and cultivated maize.

Allier, A., Teyssèdre, S., Lehermeier, C., Claustres, B., Maltese, S., Melkior, S., Moreau, L., & Charcosset, A. (2019). Assessment of breeding programs sustainability : Application of phenotypic and genomic indicators to a North European grain maize program. Theoretical and Applied Genetics, 132(5), 1321□1334. 10.1007/s00122-019-03280-w

Arca, M., Gouesnard, B., Mary□Huard, T., Le Paslier, M., Bauland, C., Combes, V., Madur, D., Charcosset, A., & Nicolas, S. D. (2023). Genotyping of DNA pools identifies untapped landraces and genomic regions to develop next generation varieties. Plant Biotechnology Journal, pbi.14022. 10.1111/pbi.14022

Arca, M., Mary-Huard, T., Gouesnard, B., Bérard, A., Bauland, C., Combes, V., Madur, D., Charcosset, A., & Nicolas, S. D. (2021). Deciphering the Genetic Diversity of Landraces With High-Throughput SNP Genotyping of DNA Bulks : Methodology and Application to the Maize 50k Array. Frontiers in Plant Science, 11, 568699. 10.3389/fpls.2020.568699

Ashraf, B. H., Jensen, J., Asp, T., & Janss, L. L. (2014). Association studies using family pools of outcrossing crops based on allele-frequency estimates from DNA sequencing. Theoretical and Applied Genetics, 127(6), 1331□1341. 10.1007/s00122-014-2300-4

Balconi, C., Galaretto, A., Malvar, R. A., Nicolas, S. D., Redaelli, R., Andjelkovic, V., Revilla, P., Bauland, C., Gouesnard, B., Butron, A., Torri, A., Barata, A. M., Kravic, N., Combes, V., Mendes-Moreira, P., Murariu, D., Šarčević, H., Schierscher-Viret, B., Vincent, M., … Goritschnig, S. (2024). Genetic and Phenotypic Evaluation of European Maize Landraces as a Tool for Conservation and Valorization of Agrobiodiversity. Biology, 13(6), 454. 10.3390/biology13060454

Beck, S. V., May, S. A., Kess, T., Bradbury, I. R., Lozada□Soto, E. A., & Wellenreuther, M. (2025). Applying Genomic Offsets to Breeding Programmes : Bridging Evolutionary Insights With Practical Applications. Evolutionary Applications, 18(10), e70155. 10.1111/eva.70155

Bernardo, R. (1994). Prediction of Maize Single□Cross Performance Using RFLPs and Information from Related Hybrids. Crop Science, 34(1), 20□25. 10.2135/cropsci1994.0011183X003400010003x

Böhm, J., Schipprack, W., Utz, H. F., & Melchinger, A. E. (2017). Tapping the genetic diversity of landraces in allogamous crops with doubled haploid lines : A case study from European flint maize. Theoretical and Applied Genetics, 130(5), 861□873. 10.1007/s00122-017-2856-x

Bohra, A., Kilian, B., Sivasankar, S., Caccamo, M., Mba, C., McCouch, S. R., & Varshney, R. K. (2022). Reap the crop wild relatives for breeding future crops. Trends in Biotechnology, 40(4), 412□431. 10.1016/j.tibtech.2021.08.009

Brandenburg, J.-T., Mary-Huard, T., Rigaill, G., Hearne, S. J., Corti, H., Joets, J., Vitte, C., Charcosset, A., Nicolas, S. D., & Tenaillon, M. I. (2017). Independent introductions and admixtures have contributed to adaptation of European maize and its American counterparts. PLOS Genetics, 13(3), e1006666. 10.1371/journal.pgen.1006666

Burgueño, J., De Los Campos, G., Weigel, K., & Crossa, J. (2012a). Genomic Prediction of Breeding Values when Modeling Genotype × Environment Interaction using Pedigree and Dense Molecular Markers. Crop Science, 52(2), 707□719. 10.2135/cropsci2011.06.0299

Burgueño, J., De Los Campos, G., Weigel, K., & Crossa, J. (2012b). Genomic Prediction of Breeding Values when Modeling Genotype × Environment Interaction using Pedigree and Dense Molecular Markers. Crop Science, 52(2), 707□719. 10.2135/cropsci2011.06.0299

Campos, H., Cooper, M., Edmeades, G. O., Loffler, C., Schussler, J. R., & Ibanez, M. (2006). Changes in drought tolerance in maize associated with fifty years of breeding for yield in the US corn belt. MAYDICA, 51(2), Article 2.

Capblancq, T., Fitzpatrick, M. C., Bay, R. A., Exposito-Alonso, M., & Keller, S. R. (2020). Genomic Prediction of (Mal)Adaptation Across Current and Future Climatic Landscapes. Annual Review of Ecology, Evolution, and Systematics, 51(1), 245□269. 10.1146/annurev-ecolsys-020720-042553

Ceccarelli, S., Grando, S., Maatougui, M., Michael, M., Slash, M., Haghparast, R., Rahmanian, M., Taheri, A., Al-Yassin, A., Benbelkacem, A., Labdi, M., Mimoun, H., & Nachit, M. (2010). Plant breeding and climate changes. The Journal of Agricultural Science, 148(6), 627□637. 10.1017/S0021859610000651

Cericola, F., Lenk, I., Fè, D., Byrne, S., Jensen, C. S., Pedersen, M. G., Asp, T., Jensen, J., & Janss, L. (2018). Optimized Use of Low-Depth Genotyping-by-Sequencing for Genomic Prediction Among Multi-Parental Family Pools and Single Plants in Perennial Ryegrass (Lolium perenne L.). Frontiers in Plant Science, 9, 369. 10.3389/fpls.2018.00369

Challinor, A. J., Koehler, A.-K., Ramirez-Villegas, J., Whitfield, S., & Das, B. (2016). Current warming will reduce yields unless maize breeding and seed systems adapt immediately. Nature Climate Change, 6(10), 954□958. 10.1038/nclimate3061

Challinor, A. J., Watson, J., Lobell, D. B., Howden, S. M., Smith, D. R., & Chhetri, N. (2014). A meta-analysis of crop yield under climate change and adaptation. Nature Climate Change, 4(4), 287□291. 10.1038/nclimate2153

Covarrubias-Pazaran, G. (2016). sommer : Solving Mixed Model Equations in R (p. 4.4.3) [Jeu de données]. 10.32614/CRAN.package.sommer

Crow, J. F., & Kimura, M. (1970). An Introduction to Population Genetics Theory. Harper & Row. https://books.google.fr/books?id=ytMPAQAAMAAJ

Duvick, D. N. (2001). Biotechnology in the 1930s : The development of hybrid maize. Nature Reviews Genetics, 2(1), 69□74. 10.1038/35047587

El Hanafi, S., Jiang, Y., Kehel, Z., Schulthess, A. W., Zhao, Y., Mascher, M., Haupt, M., Himmelbach, A., Stein, N., Amri, A., & Reif, J. C. (2023). Genomic predictions to leverage phenotypic data across genebanks. Frontiers in Plant Science, 14, 1227656. 10.3389/fpls.2023.1227656

Ellis, N., Smith, S. J., & Pitcher, C. R. (2012). Gradient forests : Calculating importance gradients on physical predictors. Ecology, 93(1), 156□168. 10.1890/11-0252.1

Fick, S. E., & Hijmans, R. J. (2017). WorldClim 2 : New 1□km spatial resolution climate surfaces for global land areas. International Journal of Climatology, 37(12), 4302□4315. 10.1002/joc.5086

Fitzpatrick, M. C., Chhatre, V. E., Soolanayakanahally, R. Y., & Keller, S. R. (2021). Experimental support for genomic prediction of climate maladaptation using the machine learning approach Gradient Forests. Molecular Ecology Resources, 21(8), 2749□2765. 10.1111/1755-0998.13374

Fitzpatrick, M. C., & Keller, S. R. (2015). Ecological genomics meets community-level modelling of biodiversity : Mapping the genomic landscape of current and future environmental adaptation. Ecology Letters, 18(1), 1□16. 10.1111/ele.12376

Fitzpatrick, M. C., Keller, S. R., & Lotterhos, K. E. (2025). The challenge of genomic forecasting in an era of global change. The American Naturalist, 738891. 10.1086/738891

Ganal, M. W., Durstewitz, G., Polley, A., Bérard, A., Buckler, E. S., Charcosset, A., Clarke, J. D., Graner, E.-M., Hansen, M., Joets, J., Le Paslier, M.-C., McMullen, M. D., Montalent, P., Rose, M., Schön, C.-C., Sun, Q., Walter, H., Martin, O. C., & Falque, M. (2011). A Large Maize (Zea mays L.) SNP Genotyping Array : Development and Germplasm Genotyping, and Genetic Mapping to Compare with the B73 Reference Genome. PLoS ONE, 6(12), e28334. 10.1371/journal.pone.0028334

Gholami, M., Wimmer, V., Sansaloni, C., Petroli, C., Hearne, S. J., Covarrubias-Pazaran, G., Rensing, S., Heise, J., Pérez-Rodríguez, P., Dreisigacker, S., Crossa, J., & Martini, J. W. R. (2021). A Comparison of the Adoption of Genomic Selection Across Different Breeding Institutions. Frontiers in Plant Science, 12, 728567. 10.3389/fpls.2021.728567

Godfray, H. C. J., Beddington, J. R., Crute, I. R., Haddad, L., Lawrence, D., Muir, J. F., Pretty, J., Robinson, S., Thomas, S. M., & Toulmin, C. (2010). Food Security : The Challenge of Feeding 9 Billion People. Science, 327(5967), 812□818. 10.1126/science.1185383

Gorjanc, G., Jenko, J., Hearne, S. J., & Hickey, J. M. (2016). Initiating maize pre-breeding programs using genomic selection to harness polygenic variation from landrace populations. BMC Genomics, 17(1), 30. 10.1186/s12864-015-2345-z

Gouesnard, B., Dallard, J., Panouillé, A., & Boyat, A. (1997). Classification of French maize populations based on morphological traits. Agronomie, 17(9□10), 491□498. 10.1051/agro:19970906

Gougherty, A. V., Keller, S. R., & Fitzpatrick, M. C. (2021). Maladaptation, migration and extirpation fuel climate change risk in a forest tree species. Nature Climate Change, 11(2), 166□171. 10.1038/s41558-020-00968-6

Guo, X., Cericola, F., Fè, D., Pedersen, M. G., Lenk, I., Jensen, C. S., Jensen, J., & Janss, L. L. (2018). Genomic Prediction in Tetraploid Ryegrass Using Allele Frequencies Based on Genotyping by Sequencing. Frontiers in Plant Science, 9, 1165. 10.3389/fpls.2018.01165

Hallauer, A. R., Carena, M. J., & Miranda Filho, J. B. (2010). Quantitative genetics in maize breeding (3rd ed.). Springer.

Hawkesford, M. J., & Griffiths, S. (2019). Exploiting genetic variation in nitrogen use efficiency for cereal crop improvement. Current Opinion in Plant Biology, 49, 35□42. 10.1016/j.pbi.2019.05.003

Heino, M., Kinnunen, P., Anderson, W., Ray, D. K., Puma, M. J., Varis, O., Siebert, S., & Kummu, M. (2023). Increased probability of hot and dry weather extremes during the growing season threatens global crop yields. Scientific Reports, 13(1), 3583. 10.1038/s41598-023-29378-2

Hijmans, R. J. (2010). raster : Geographic Data Analysis and Modeling (p. 3. 6-32) [Jeu de données]. 10.32614/CRAN.package.raster

Hijmans, R. J. (2020). terra : Spatial Data Analysis (p. 1. 8-70) [Jeu de données]. 10.32614/CRAN.package.terra

Hijmans, R. J., Phillips, S., Leathwick, J., & Elith, J. (2010). dismo : Species Distribution Modeling (p. 1. 3-16) [Jeu de données]. 10.32614/CRAN.package.dismo

Hoisington, D., Khairallah, M., Reeves, T., Ribaut, J.-M., Skovmand, B., Taba, S., & Warburton, M. (1999). Plant genetic resources : What can they contribute toward increased crop productivity? Proceedings of the National Academy of Sciences, 96(11), 5937□5943. 10.1073/pnas.96.11.5937

Hölker, A. C., Mayer, M., Presterl, T., Bauer, E., Ouzunova, M., Melchinger, A. E., & Schön, C.-C. (2022). Theoretical and experimental assessment of genome-based prediction in landraces of allogamous crops. Proceedings of the National Academy of Sciences, 119(18), e2121797119. 10.1073/pnas.2121797119

Hölker, A. C., Mayer, M., Presterl, T., Bolduan, T., Bauer, E., Ordas, B., Brauner, P. C., Ouzunova, M., Melchinger, A. E., & Schön, C.-C. (2019). European maize landraces made accessible for plant breeding and genome-based studies. Theoretical and Applied Genetics, 132(12), 3333□3345. 10.1007/s00122-019-03428-8

Hudson, A. I., Odell, S. G., Dubreuil, P., Tixier, M.-H., Praud, S., Runcie, D. E., & Ross-Ibarra, J. (2022). Analysis of genotype-by-environment interactions in a maize mapping population. G3 Genes|Genomes|Genetics, 12(3), jkac013. 10.1093/g3journal/jkac013

Jägermeyr, J., Müller, C., Ruane, A. C., Elliott, J., Balkovic, J., Castillo, O., Faye, B., Foster, I., Folberth, C., Franke, J. A., Fuchs, K., Guarin, J. R., Heinke, J., Hoogenboom, G., Iizumi, T., Jain, A. K., Kelly, D., Khabarov, N., Lange, S., … Rosenzweig, C. (2021). Climate impacts on global agriculture emerge earlier in new generation of climate and crop models. Nature Food, 2(11), 873□885. 10.1038/s43016-021-00400-y

Jarquín, D., Lemes Da Silva, C., Gaynor, R. C., Poland, J., Fritz, A., Howard, R., Battenfield, S., & Crossa, J. (2017). Increasing Genomic□Enabled Prediction Accuracy by Modeling Genotype × Environment Interactions in Kansas Wheat. The Plant Genome, 10(2), plantgenome2016.12.0130. 10.3835/plantgenome2016.12.0130

Keep, T., Rouet, S., Blanco-Pastor, J. L., Barre, P., Ruttink, T., Dehmer, K. J., Hegarty, M., Ledauphin, T., Litrico, I., Muylle, H., Roldán-Ruiz, I., Surault, F., Veron, R., Willner, E., & Sampoux, J. P. (2021). Inter-annual and spatial climatic variability have led to a balance between local fluctuating selection and wide-range directional selection in a perennial grass species. Annals of Botany, 128(3), 357□369. 10.1093/aob/mcab057

Keep, T., Sampoux, J.-P., Blanco-Pastor, J. L., Dehmer, K. J., Hegarty, M. J., Ledauphin, T., Litrico, I., Muylle, H., Roldán-Ruiz, I., Roschanski, A. M., Ruttink, T., Surault, F., Willner, E., & Barre, P. (2020). High-Throughput Genome-Wide Genotyping To Optimize the Use of Natural Genetic Resources in the Grassland Species Perennial Ryegrass (Lolium perenne L.). G3 Genes|Genomes|Genetics, 10(9), 3347□3364. 10.1534/g3.120.401491

Knox, J., Daccache, A., Hess, T., & Haro, D. (2016). Meta-analysis of climate impacts and uncertainty on crop yields in Europe. Environmental Research Letters, 11(11), 113004. 10.1088/1748-9326/11/11/113004

Kotni, P., van Hintum, T., Maggioni, L., Oppermann, M., & Weise, S. (2023). EURISCO update 2023 : The European Search Catalogue for Plant Genetic Resources, a pillar for documentation of genebank material. Nucleic Acids Research, 51(D1), D1465□D1469. 10.1093/nar/gkac852

Li, F., Gates, D. J., Buckler, E. S., Hufford, M. B., Janzen, G. M., Rellán-Álvarez, R., Rodríguez-Zapata, F., Romero Navarro, J. A., Sawers, R. J. H., Snodgrass, S. J., Sonder, K., Willcox, M. C., Hearne, S. J., Ross-Ibarra, J., & Runcie, D. E. (2025). Environmental data provide marginal benefit for predicting climate adaptation. PLOS Genetics, 21(6), e1011714. 10.1371/journal.pgen.1011714

Lotterhos, K. E. (2024). Interpretation issues with “genomic vulnerability” arise from conceptual issues in local adaptation and maladaptation. Evolution Letters, 8(3), 331□339. 10.1093/evlett/qrae004

Martini, J. W. R., Molnar, T. L., Crossa, J., Hearne, S. J., & Pixley, K. V. (2021). Opportunities and Challenges of Predictive Approaches for Harnessing the Potential of Genetic Resources. Frontiers in Plant Science, 12, 674036. 10.3389/fpls.2021.674036

Mascher, M., Schreiber, M., Scholz, U., Graner, A., Reif, J. C., & Stein, N. (2019). Genebank genomics bridges the gap between the conservation of crop diversity and plant breeding. Nature Genetics, 51(7), 1076□1081. 10.1038/s41588-019-0443-6

Matsuoka, Y., Vigouroux, Y., Goodman, M. M., Sanchez G. J., Buckler, E., & Doebley, J. (2002). A single domestication for maize shown by multilocus microsatellite genotyping. Proceedings of the National Academy of Sciences, 99(9), 6080□6084. 10.1073/pnas.052125199

Mayer, M., Hölker, A. C., González-Segovia, E., Bauer, E., Presterl, T., Ouzunova, M., Melchinger, A. E., & Schön, C.-C. (2020). Discovery of beneficial haplotypes for complex traits in maize landraces. Nature Communications, 11(1), 4954. 10.1038/s41467-020-18683-3

McLaughlin, C. M., Shi, Y., Viswanathan, V., Sawers, R., Kemanian, A. R., & Lasky, J. R. (2024). Maladaptation in cereal crop landraces following a soot-producing climate catastrophe. 10.1101/2024.05.18.594591

Meuwissen, T. H. E., Hayes, B. J., & Goddard, M. E. (2001). Prediction of Total Genetic Value Using Genome-Wide Dense Marker Maps. Genetics, 157(4), 1819□1829. 10.1093/genetics/157.4.1819

Minoli, S., Jägermeyr, J., Asseng, S., Urfels, A., & Müller, C. (2022). Global crop yields can be lifted by timely adaptation of growing periods to climate change. Nature Communications, 13(1), 7079. 10.1038/s41467-022-34411-5

Mir, C., Zerjal, T., Combes, V., Dumas, F., Madur, D., Bedoya, C., Dreisigacker, S., Franco, J., Grudloyma, P., Hao, P. X., Hearne, S., Jampatong, C., Laloë, D., Muthamia, Z., Nguyen, T., Prasanna, B. M., Taba, S., Xie, C. X., Yunus, M., … Charcosset, A. (2013). Out of America : Tracing the genetic footprints of the global diffusion of maize. Theoretical and Applied Genetics, 126(11), 2671□2682. 10.1007/s00122-013-2164-z

Nei, M. (1973). Analysis of Gene Diversity in Subdivided Populations. Proceedings of the National Academy of Sciences, 70(12), 3321□3323. 10.1073/pnas.70.12.3321

OECD & Food and Agriculture Organization of the United Nations. (2022). OECD-FAO Agricultural Outlook 2022-2031. OECD. 10.1787/f1b0b29c-en

Parent, B., Leclere, M., Lacube, S., Semenov, M. A., Welcker, C., Martre, P., & Tardieu, F. (2018). Maize yields over Europe may increase in spite of climate change, with an appropriate use of the genetic variability of flowering time. Proceedings of the National Academy of Sciences, 115(42), 10642□10647. 10.1073/pnas.1720716115

Pégard, M., Barre, P., Delaunay, S., Surault, F., Karagić, D., Milić, D., Zorić, M., Ruttink, T., & Julier, B. (2023). Genome-wide genotyping data renew knowledge on genetic diversity of a worldwide alfalfa collection and give insights on genetic control of phenology traits. Frontiers in Plant Science, 14, 1196134. 10.3389/fpls.2023.1196134

Rebourg, C., Chastanet, M., Gouesnard, B., Welcker, C., Dubreuil, P., & Charcosset, A. (2003). Maize introduction into Europe : The history reviewed in the light of molecular data. Theoretical and Applied Genetics, 106(5), 895□903. 10.1007/s00122-002-1140-9

Rebourg, C., Gouesnard, B., & Charcosset, A. (2001). Large scale molecular analysis of traditional European maize populations. Relationships with morphological variation. Heredity, 86(5), 574□587. 10.1046/j.1365-2540.2001.00869.x

Redaelli, R., Bassolino, L., Balconi, C., Terracciano, I., Torri, A., Nicoletti, F., Benedetti, G., Iacoponi, V., Rea, R., & Taviani, P. (2024). Morpho-Phenological, Chemical, and Genetic Characterization of Italian Maize Landraces from the Lazio Region. Plants, 13(22), 3249. 10.3390/plants13223249

Rellstab, C., Dauphin, B., & Exposito□Alonso, M. (2021). Prospects and limitations of genomic offset in conservation management. Evolutionary Applications, 14(5), 1202□1212. 10.1111/eva.13205

Rezaei, E. E., Webber, H., Asseng, S., Boote, K., Durand, J. L., Ewert, F., Martre, P., & MacCarthy, D. S. (2023). Climate change impacts on crop yields. Nature Reviews Earth & Environment, 4(12), 831□846. 10.1038/s43017-023-00491-0

Rhoné, B., Defrance, D., Berthouly-Salazar, C., Mariac, C., Cubry, P., Couderc, M., Dequincey, A., Assoumanne, A., Kane, N. A., Sultan, B., Barnaud, A., & Vigouroux, Y. (2020). Pearl millet genomic vulnerability to climate change in West Africa highlights the need for regional collaboration. Nature Communications, 11(1), 5274. 10.1038/s41467-020-19066-4

Rincent, R., Laloë, D., Nicolas, S., Altmann, T., Brunel, D., Revilla, P., Rodríguez, V. M., Moreno-Gonzalez, J., Melchinger, A., Bauer, E., Schoen, C.-C., Meyer, N., Giauffret, C., Bauland, C., Jamin, P., Laborde, J., Monod, H., Flament, P., Charcosset, A., & Moreau, L. (2012). Maximizing the Reliability of Genomic Selection by Optimizing the Calibration Set of Reference Individuals : Comparison of Methods in Two Diverse Groups of Maize Inbreds (Zea mays L.). Genetics, 192(2), 715□728. 10.1534/genetics.112.141473

Rogers, A. R., Dunne, J. C., Romay, C., Bohn, M., Buckler, E. S., Ciampitti, I. A., Edwards, J., Ertl, D., Flint-Garcia, S., Gore, M. A., Graham, C., Hirsch, C. N., Hood, E., Hooker, D. C., Knoll, J., Lee, E. C., Lorenz, A., Lynch, J. P., McKay, J., … Holland, J. B. (2021). The importance of dominance and genotype-by-environment interactions on grain yield variation in a large-scale public cooperative maize experiment. G3 Genes|Genomes|Genetics, 11(2), jkaa050. 10.1093/g3journal/jkaa050

Romero Navarro, J. A., Willcox, M., Burgueño, J., Romay, C., Swarts, K., Trachsel, S., Preciado, E., Terron, A., Delgado, H. V., Vidal, V., Ortega, A., Banda, A. E., Montiel, N. O. G., Ortiz-Monasterio, I., Vicente, F. S., Espinoza, A. G., Atlin, G., Wenzl, P., Hearne, S., & Buckler, E. S. (2017). A study of allelic diversity underlying flowering-time adaptation in maize landraces. Nature Genetics, 49(3), 476□480. 10.1038/ng.3784

Roth, M., Beugnot, A., Mary-Huard, T., Moreau, L., Charcosset, A., & Fiévet, J. B. (2022). Improving genomic predictions with inbreeding and nonadditive effects in two admixed maize hybrid populations in single and multienvironment contexts. Genetics, 220(4), iyac018. 10.1093/genetics/iyac018

Ruiz Corral, J. A., Durán Puga, N., Sánchez González, J. D. J., Ron Parra, J., González Eguiarte, D. R., Holland, J. B., & Medina García, G. (2008). Climatic Adaptation and Ecological Descriptors of 42 Mexican Maize Races. Crop Science, 48(4), 1502□1512. 10.2135/cropsci2007.09.0518

Sanchez, D., Sadoun, S. B., Mary-Huard, T., Allier, A., Moreau, L., & Charcosset, A. (2023). Improving the use of plant genetic resources to sustain breeding programs’ efficiency. Proceedings of the National Academy of Sciences, 120(14), e2205780119. 10.1073/pnas.2205780119

Smith, J. S., Trevisan, W., McCunn, A., & Huffman, W. E. (2022). Global dependence on Corn Belt Dent maize germplasm : Challenges and opportunities. Crop Science, 62(6), 2039□2066. 10.1002/csc2.20802

Stenberg, J. A., & Ortiz, R. (2021). Focused Identification of Germplasm Strategy (FIGS) : Polishing a rough diamond. Current Opinion in Insect Science, 45, 1□6. 10.1016/j.cois.2020.11.001

Tamang, A., Macharia, M. W., Caproni, L., Miculan, M., Mager, S., Ahmed, J. S., Yangzome, T., Pe, M. E., & Dell’Acqua, M. (2024). Genomic, climatic, and cultural diversity of maize landraces from the Himalayan Kingdom of Bhutan. PLANTS, PEOPLE, PLANET, 6(4), 965□978. 10.1002/ppp3.10513

The Third Report on The State of the World’s Plant Genetic Resources for Food and Agriculture. (2025). FAO. 10.4060/cd4711en

Tsubo, M., & Moeletsi, M. (2025). Climate risk adaptation in dryland maize through cultivar adoption. Npj Sustainable Agriculture, 3(1), 48. 10.1038/s44264-025-00088-8

Welcker, C., Spencer, N. A., Turc, O., Granato, I., Chapuis, R., Madur, D., Beauchene, K., Gouesnard, B., Draye, X., Palaffre, C., Lorgeou, J., Melkior, S., Guillaume, C., Presterl, T., Murigneux, A., Wisser, R. J., Millet, E. J., van Eeuwijk, F., Charcosset, A., & Tardieu, F. (2022). Physiological adaptive traits are a potential allele reservoir for maize genetic progress under challenging conditions. Nature Communications, 13(1), 3225. 10.1038/s41467-022-30872-w

Wickham, H., Chang, W., Henry, L., Pedersen, T. L., Takahashi, K., Wilke, C., Woo, K., Yutani, H., Dunnington, D., & Van Den Brand, T. (2007). ggplot2 : Create Elegant Data Visualisations Using the Grammar of Graphics (p. 4.0.0) [Jeu de données]. 10.32614/CRAN.package.ggplot2

Zhao, C., Liu, B., Piao, S., Wang, X., Lobell, D. B., Huang, Y., Huang, M., Yao, Y., Bassu, S., Ciais, P., Durand, J.-L., Elliott, J., Ewert, F., Janssens, I. A., Li, T., Lin, E., Liu, Q., Martre, P., Müller, C., … Asseng, S. (2017). Temperature increase reduces global yields of major crops in four independent estimates. Proceedings of the National Academy of Sciences, 114(35), 9326□9331. 10.1073/pnas.1701762114

