## Supplementary Figures for "A new evolution-based genomic prediction model forecasts yield performance across environments and future climates and identifies adapted maize landraces"

#### Figure S1 – Gradient Forest model outputs

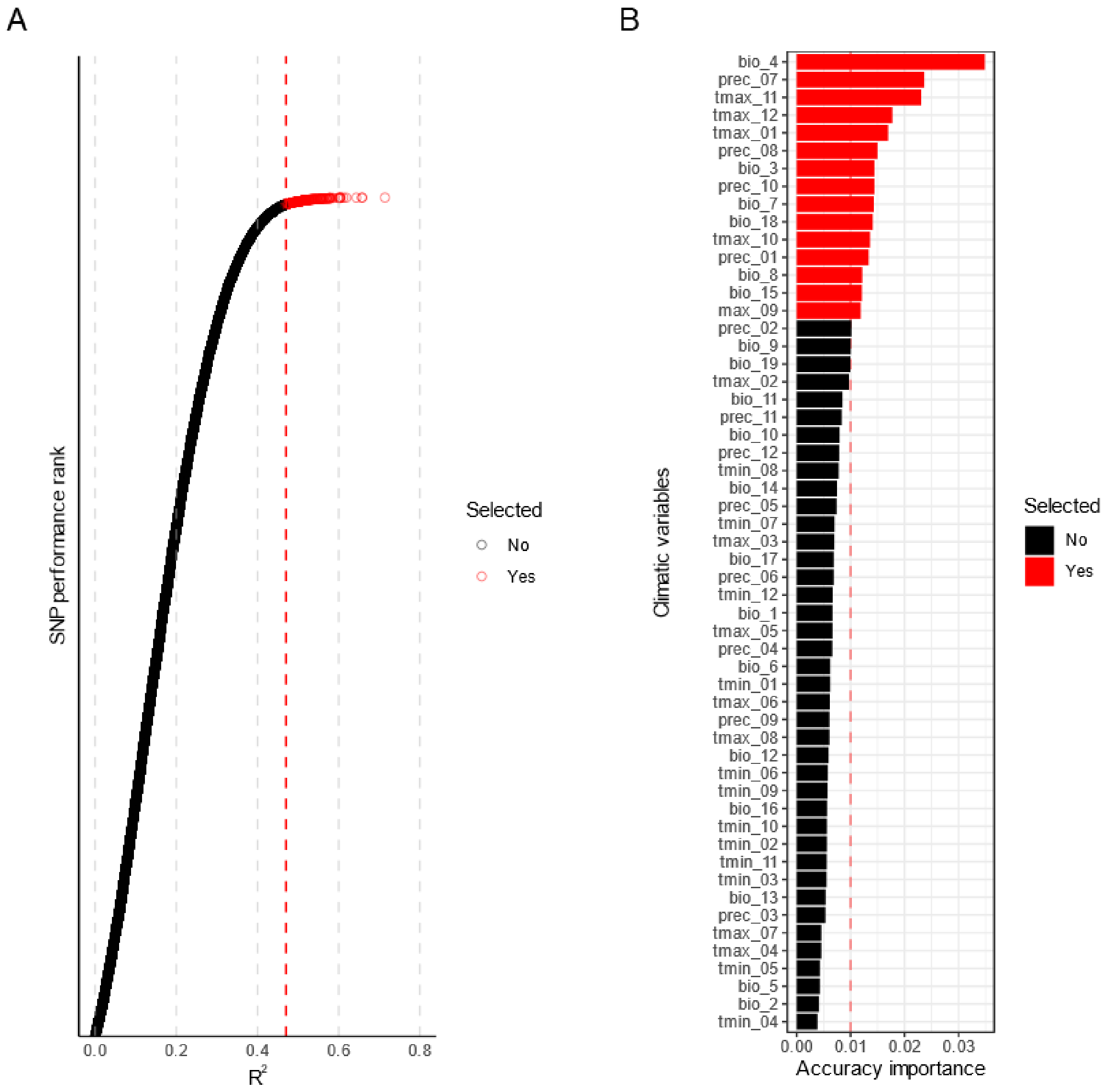

**Figure S1: Classification of SNPs and climatic variables according to their importance to predict adaptive allelic frequency variation across environment using gradient forest models.** (**A**) Gradient forest (GF) model goodness-of-fit ($R^2$) for each SNP using all climatic variables from WorldClim2.1. (**B**) Accuracy importance for the 55 climatic variables estimated using a second GF model including only 243 SNPs that were best predicted by the first GF model (highest $R^2$). SNPs (**A**) and climatic variables (**B**) selected by these two successive GF models are highlighted in red.

#### Figure S2 – Local genomic offset for SSP585 scenario at 2081-2100.

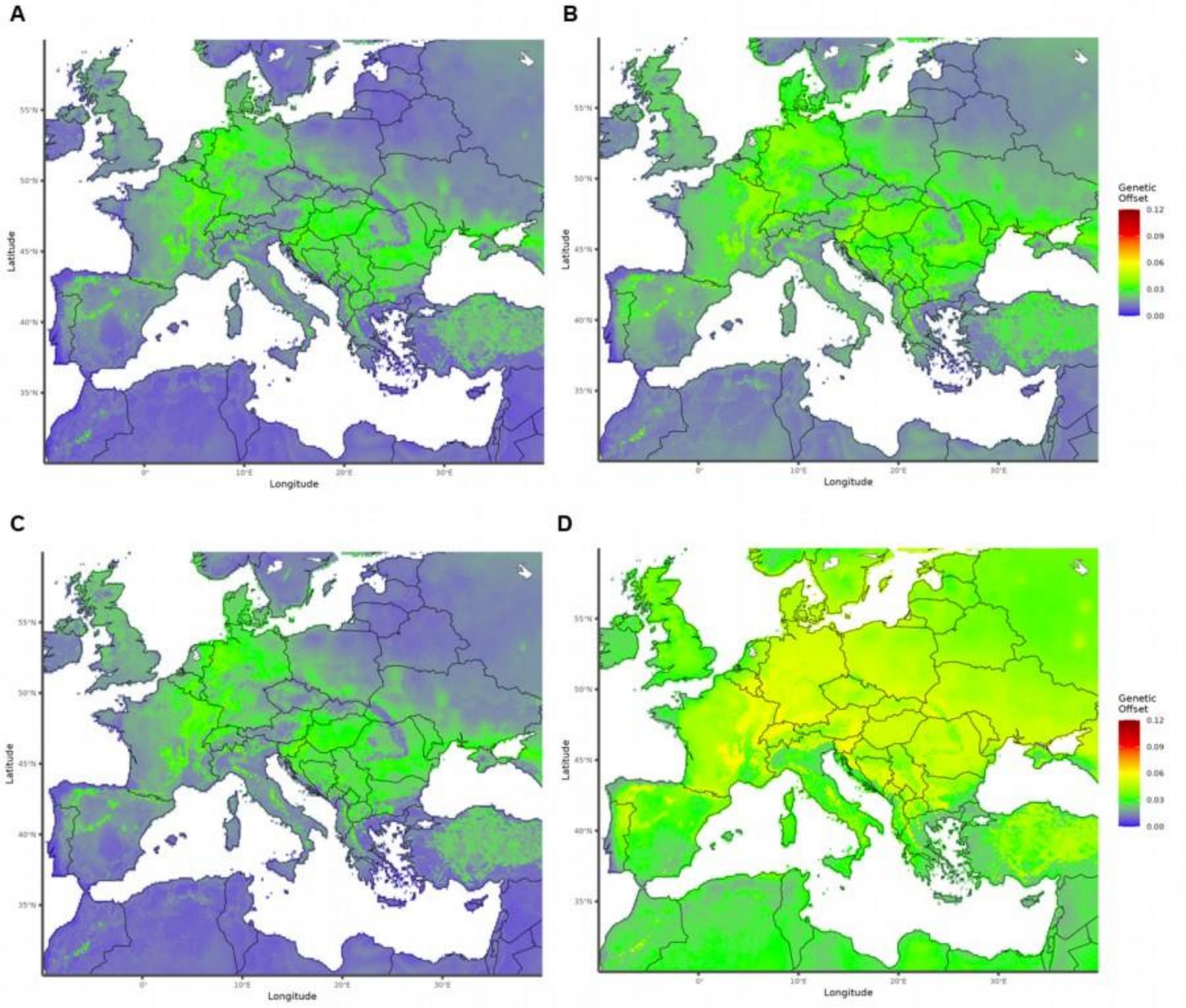

**Figure S2: Variation of local genomic offset for two future climatic scenarios**. Map including geographic distribution of local genomic offset (LGO) based on Gradient Forest model under greenhouse gases emission pathways SSP126 (**A**, **B**) and SSP585 (**C**, **D**) for 2041-2060 (**A**, **C**) and 2081-2100 (**B**, **D**) time horizons. Genomic Offset (GO) is estimated as the GO between each grid in past climatic conditions (1979-2000) and the same grid in a future climatic scenario.

### Figure S3 – Phenotypic variation per trial

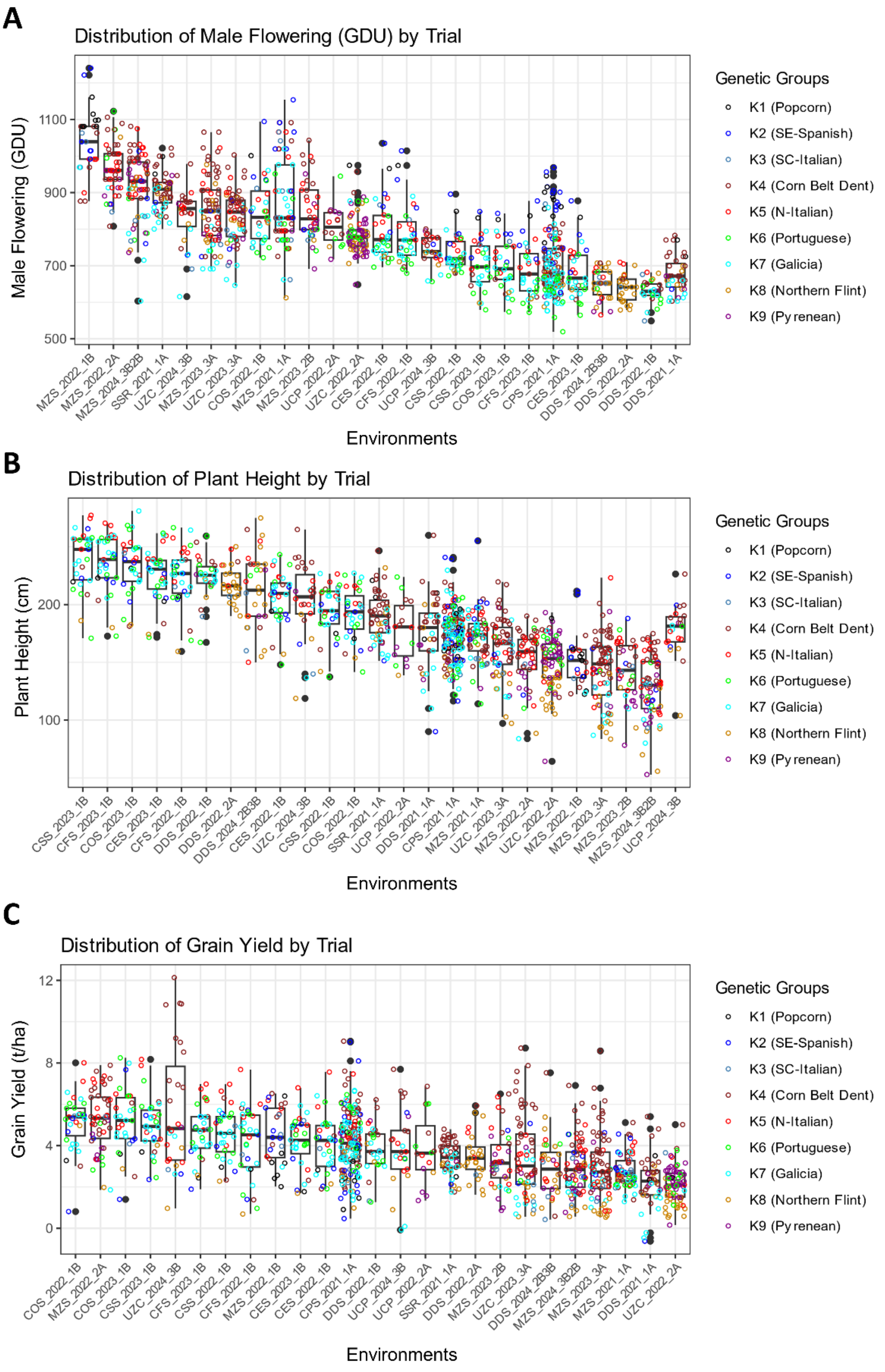

**Figure S3: Phenotypic variation in EVA Network trials**. Each boxplot represents the distribution of the phenotypic adjusted means in each environment for male flowering (**A**), plant height (**B**) and grain yield (**C**). Each point represents one landrace in one corresponding environment. Points are colored according to their maximum assignment to nine genetic groups.

#### Figure S4 – GO explored in the EVA Network per landrace

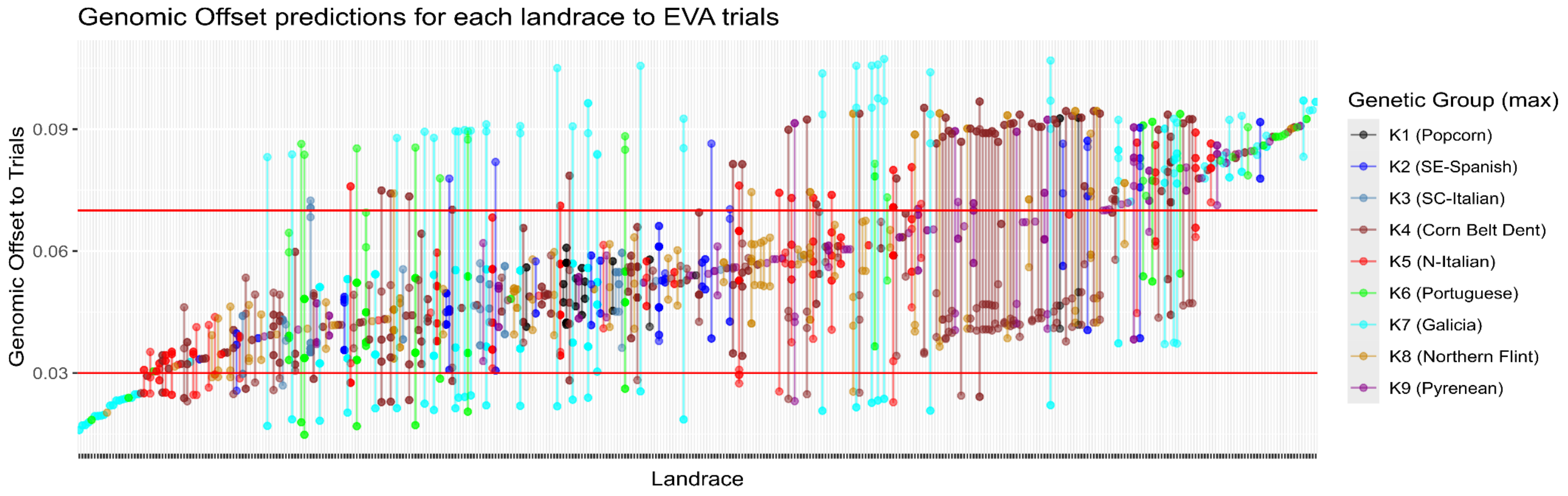

**Figure S4: Variation of genomic offset for each landrace across EVA field trial experimental network.** Each dot represents one trial-landrace combination (470 in total), and the vertical lines represent the range of genomic offset explored by each landrace in field trial experimental networks. Both the dots and the lines are colored according to the assignment of each landrace to one of the nine genetic groups considering the maximum assignment. Horizontal lines in red represent the GO limits of 0.03 and 0.07, that were considered as a reference of "low" and "high" genomic offset, respectively.

#### Figure S5 – Relationship of GO and Geographical distance to EVA Network trials

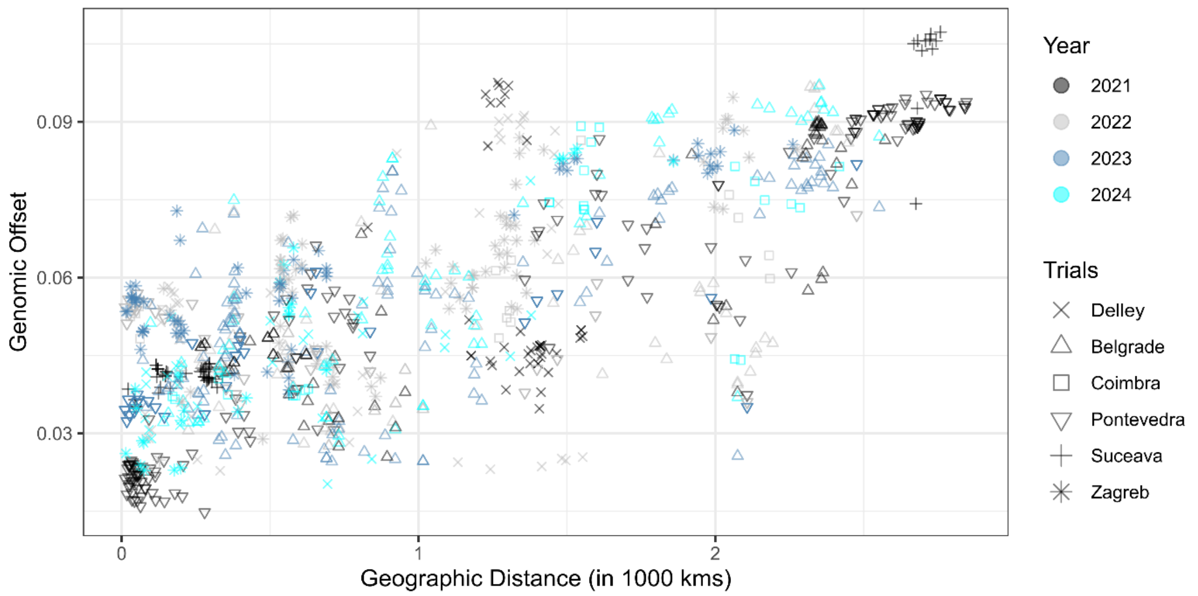

**Figure S5: Genomic offset is correlated with the geographic distance between collection sites and field experiments.** Relationship between geographic distances (thousand kilometers) from landrace collection sites to EVA Network field experiments and genomic offset. Trials sites are represented by symbols and years by colors. Each point represents one landrace evaluated in one environment (site-year combination).

#### Figure S6 – Relationship of GO with male flowering and plant height

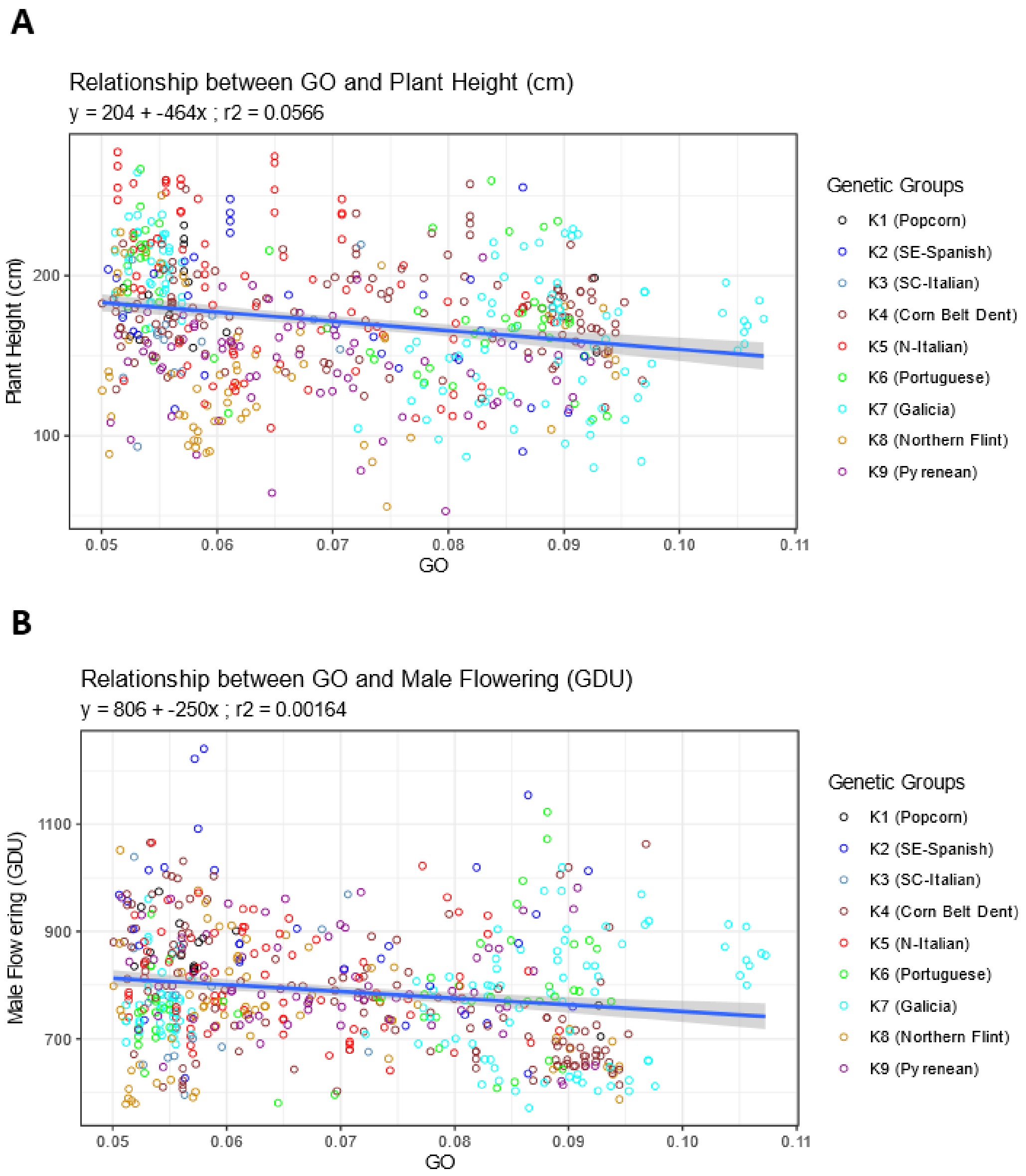

**Figure S6: Genomic offset effect on plant height and male flowering**. Relationship of genomic offset (GO) with within experiment adjusted means for plant height (PH, in cm) (**A**) and male flowering (MF, in GDU) (**B**) across the EVA Network. Each point represents one landrace in one environment, colored by their maximum assignment to genetic groups.

#### Figure S7 – Relationship of GO,for landraces evaluated under GO < 0.03 and GO > 0.07

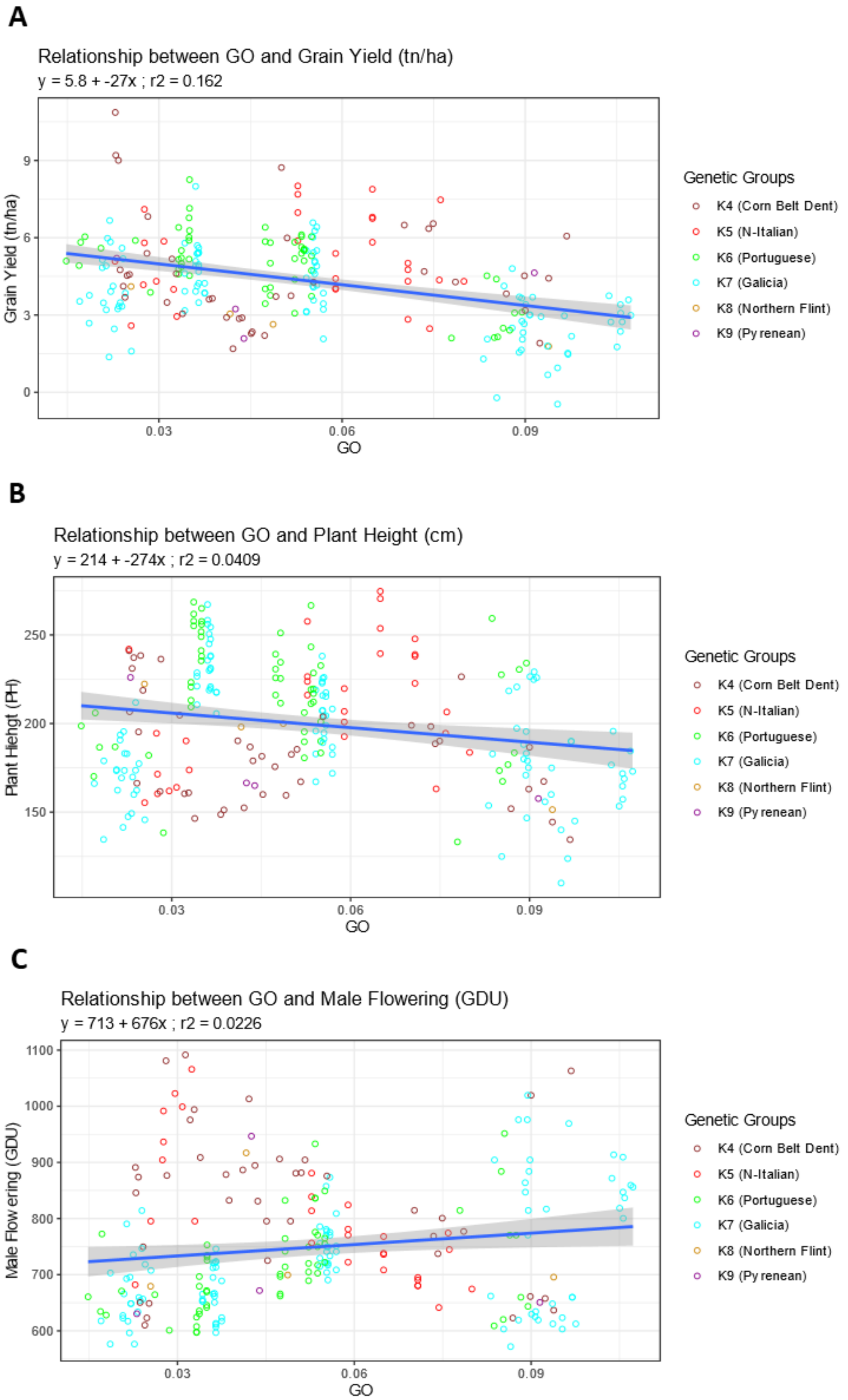

**Figure S7: Effect of genomic offset (GO) on phenotypes for landraces evaluated at high and low GO.** Relationship of genomic offset (GO) with within trial adjusted means for grain yield (GY, in tons per hectare, **A**), plant height (PH, in cm, **B**) and male flowering (MF, in GDU, **C**). Only landraces that were evaluated in environments representing high (> 0.7) and low (< 0.3) GO are included. Each point represents one landrace in one environment, colored by their maximum assignment to genetic groups.

#### Figure S8 – Relationship of Hs with male flowering and plant height

**A**

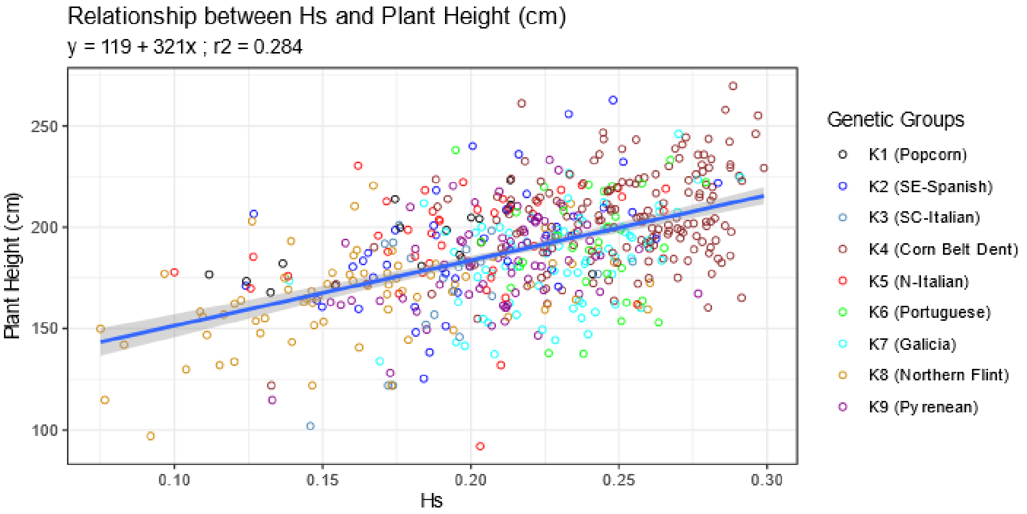

**B**

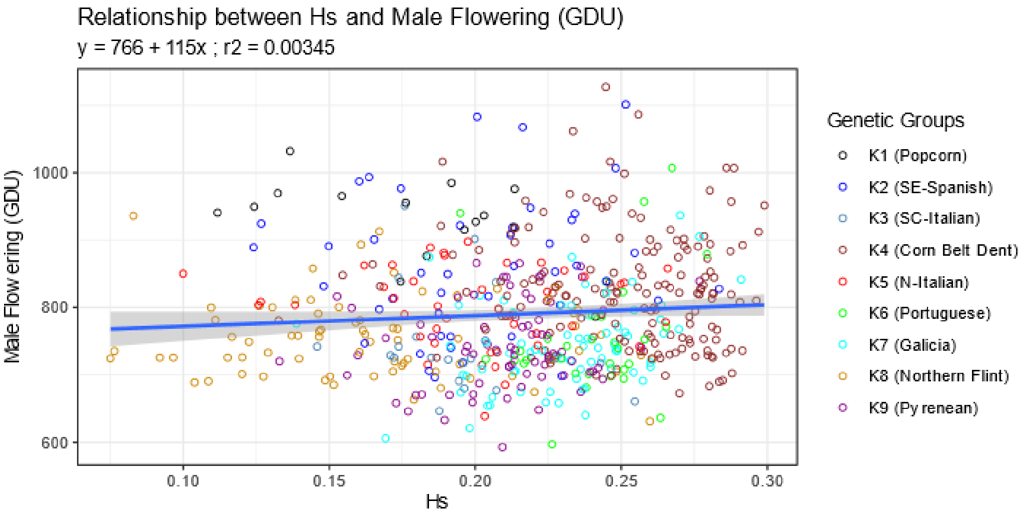

**Figure S8: Within-population gene diversity effect on plant height and male flowering.** Relationship of within population gene diversity (Hs) with across environments least square means for plant height (**A**) and male flowering (**B**). Each dot represents one landrace and is colored according to the maximum assignment to one of the nine genetic groups.

#### Figure S9 – Predicted stability vs Observed Stability and Predicted Grain Yield

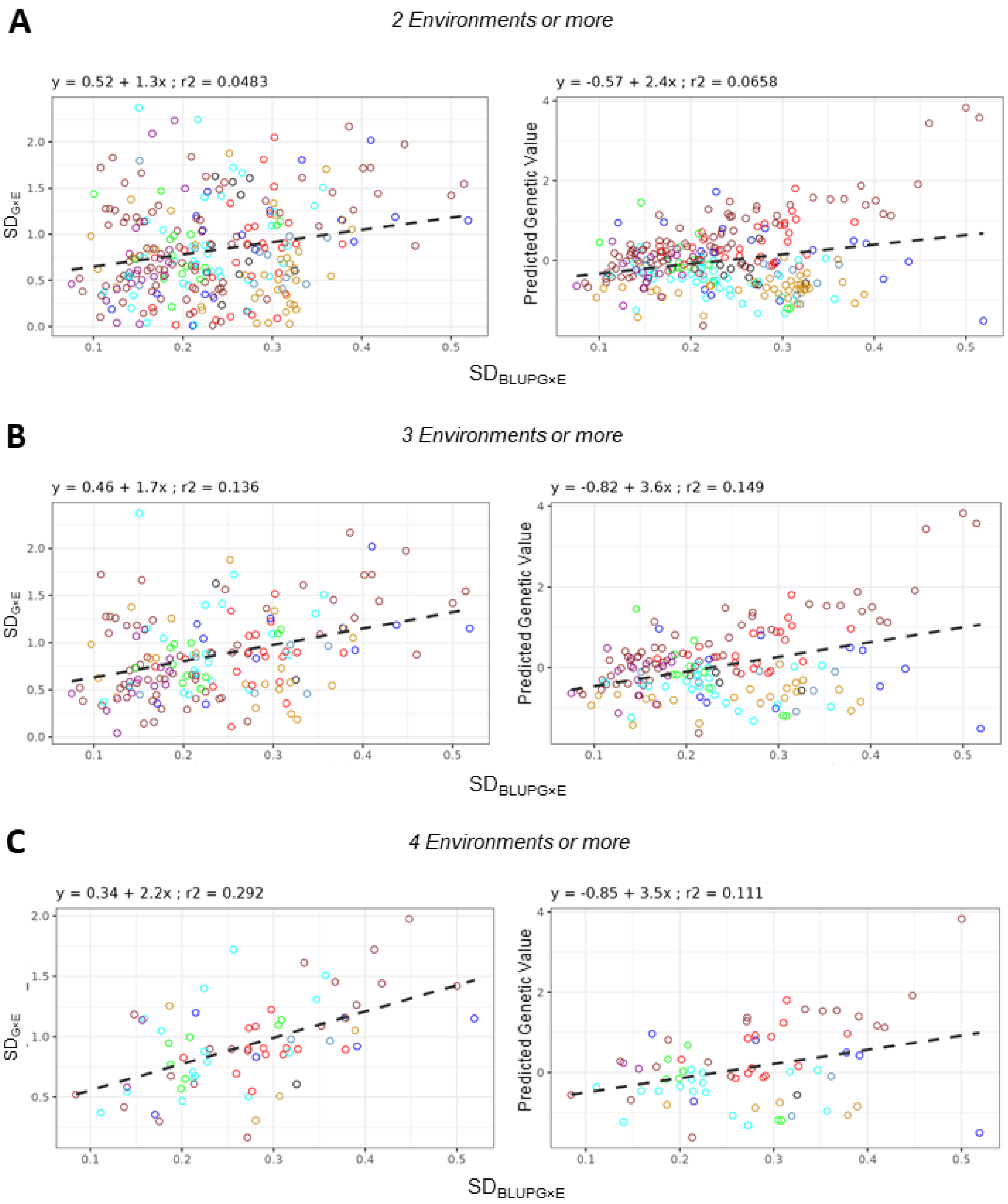

**Figure S9: Positive relationship of grain yield predicted stability with observed stability and genetic value at different levels of information.** Relationship between the grain yield predicted stability ($SD_{BLUPG \times E}$) and the observed stability ($SD_{G \times E}$) (left) and genetic value (right) for a minimum number of environments where each landrace was evaluated of two (**A**), three (**B**) or four (**C**). $SD_{G \times E}$ were obtained by first subtracting to each landrace within-environment adjusted means the least square means of the respective landrace and environment and then computing the standard deviation. The predicted stability $SD_{BLUPG \times E}$ were obtained by computing the standard deviation of genotype by environment interaction best linear unbiased predictors. The predicted genetic values are the genotypic best linear unbiased prediction from genomic prediction. Each point represents one landrace, colored by maximum assignment to genetic groups.

#### Figure S10 – Number of environments and Hs, observed and predicted yield stability.

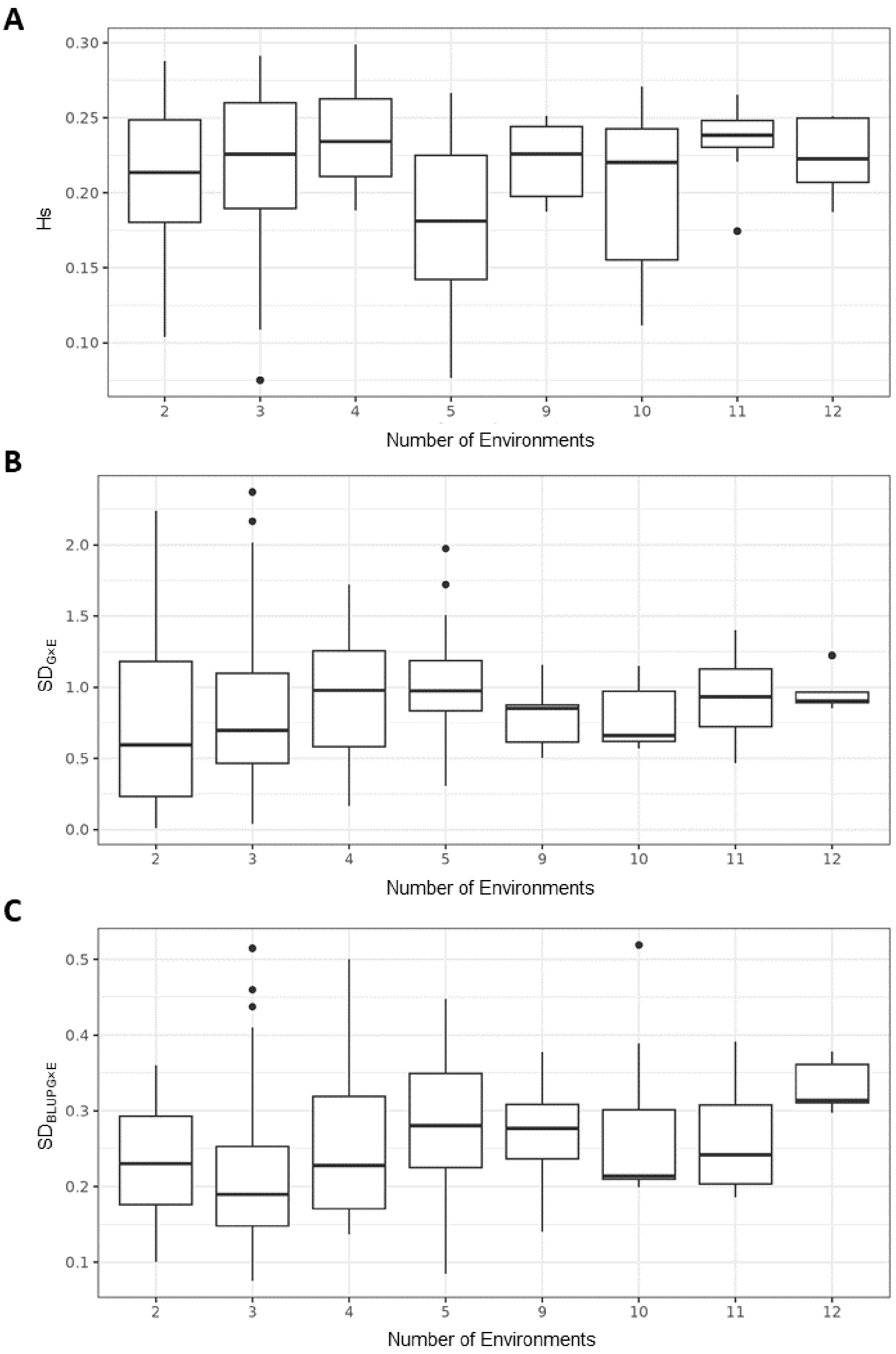

**Figure S10: Relationship between within-population gene diversity, observed, predicted $SD_{GxE}$ with the number of environments in which landraces was evaluated.** Distribution of Hs (**A**), standard deviation of observed genotype by environment interaction ($SD_{G\times E}$) (**B**) and the standard deviation of genotype by environment interaction best linear unbiased predictors ($SD_{BLUPG\times E}$) (**C**) for landraces that were evaluated at different numbers of environments.

#### Figure S11 – Cross validation schemes

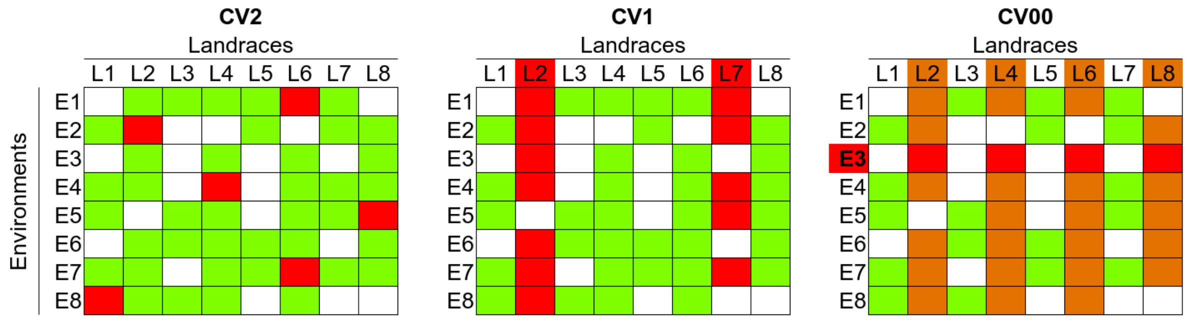

**Figure S11: Cross-validation schemes to address different prediction challenges.** Schematic representation of three different cross-validation scenarios for genomic prediction. Under **CV2 (left)**, 20% of phenotypic observations (genotype-environment combinations) were randomly hidden from the whole dataset and then predicted using 80% remaining training set. Under **CV1 (middle)**, 20% of the genotypes were randomly selected (L2 and L7 in the example) and all their phenotypic observations across all the dataset were hidden; under this scheme, genotypes to be predicted were not observed in any training dataset. Under **CV00 (right)**, each trial (E3 in the example) was hidden at each fold and the genotypes that participate in the hidden trial (L2, L4, L6 and L8 in the example) were also removed across other trials; as a consequence, neither the environment nor the genotypes predicted were observed in any training dataset. Cells in **green** represent observations that were included in the training set; cells in **red** represent observations that were removed from the data set and were then used as the validation set; cells in **orange** represent observations that were removed from the training data, but were not used for validation; cells in **white** represent genotype x environment combinations that were never observed in the network.

#### Figure S12 – Predictive Abilities comparison between M1 and Genetic Structure

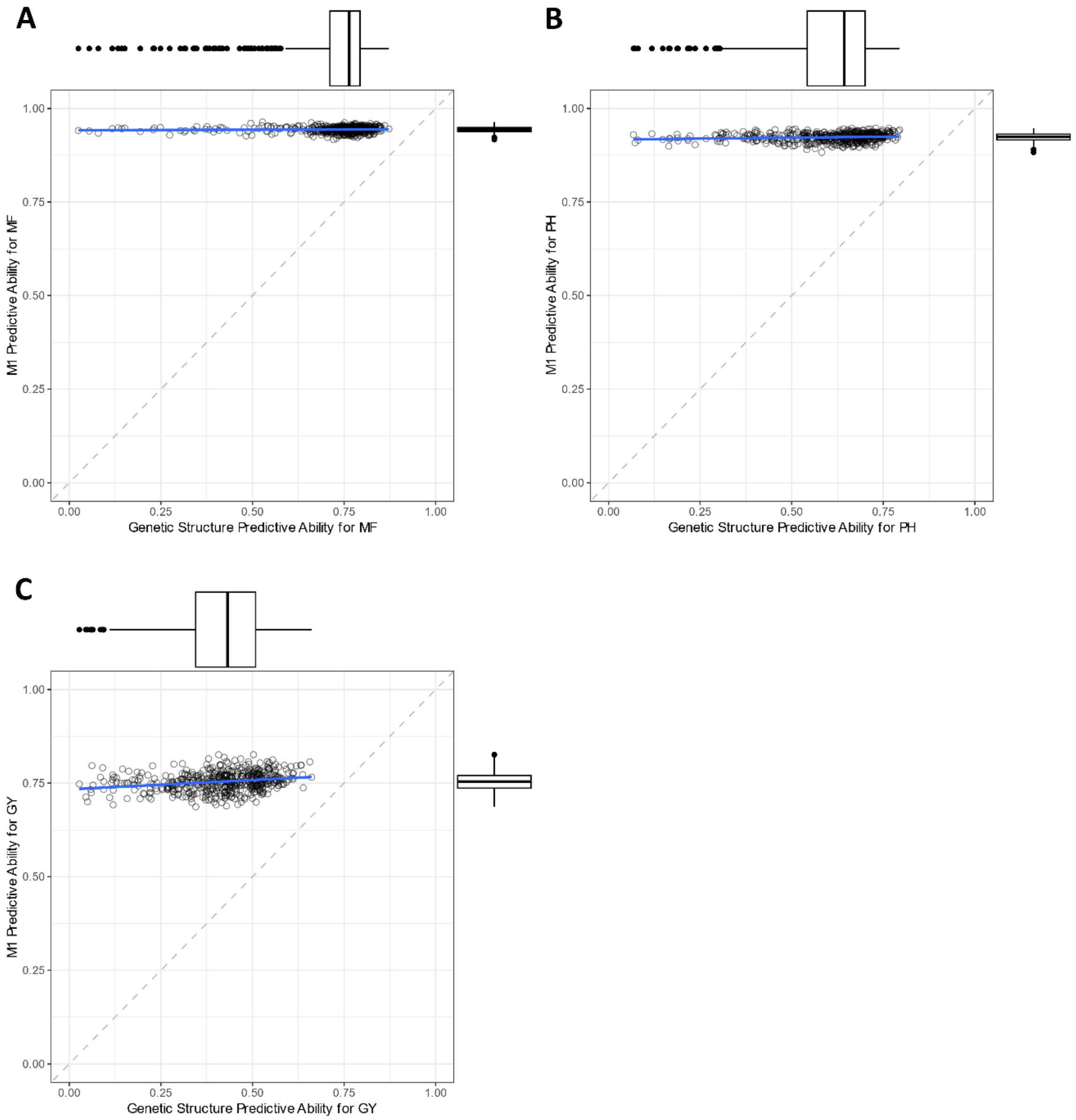

**Figure S12: Comparison of predictive ability between genomic prediction model based on the relatedness between landraces (Model 1) and based on genetic structuration.** Comparison of predictive ability obtained using Model 1 and multiple linear regression on genetic groups assignment for male flowering (MF) **(A)**, plant height (PH) **(B)** and grain yield (GY) **(C)**. Each point represents one of the five folds for each of the 100 repetitions under CV2.

Figure S13 – Predictive Ability of grain yield for each fold and model under CV00

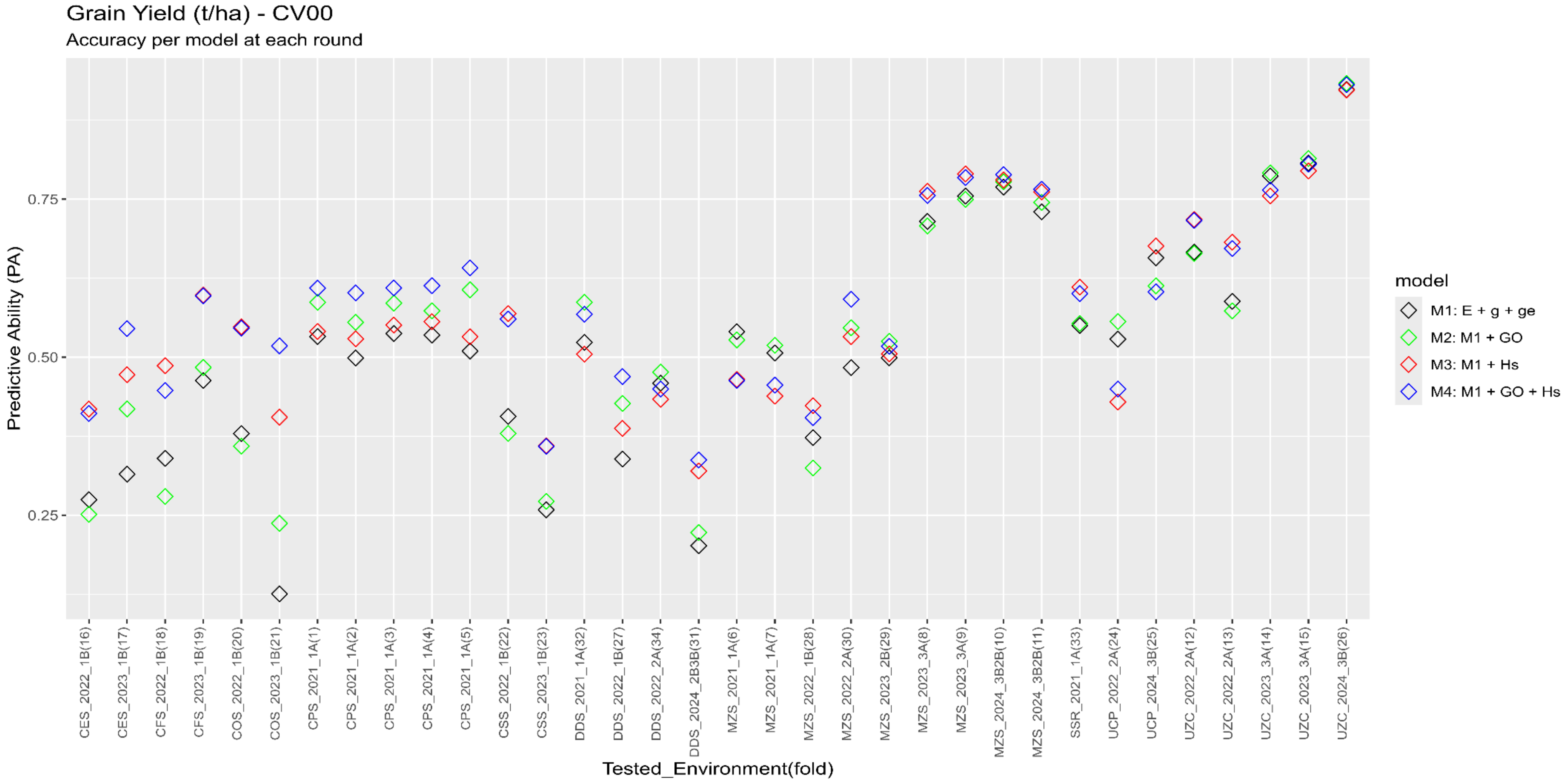

**Figure S13: Variation of predictive ability using different models when predicting each environment.** Predictive abilities of grain yield for each model and each fold of CV00 cross validation (unobserved environment and landraces). Each point represents the PA for one environment under one model. Model 1 (M1) includes random genotype and G×E effects, Model 2 (M2) adds GO fixed effect to M1, Model 3 (M3) adds Hs to M1 and Model 4 (M4) ads both GO and Hs to M1.

#### Figure S14 – Selection of interesting landraces predicted grain yield and predicted grain yield stability

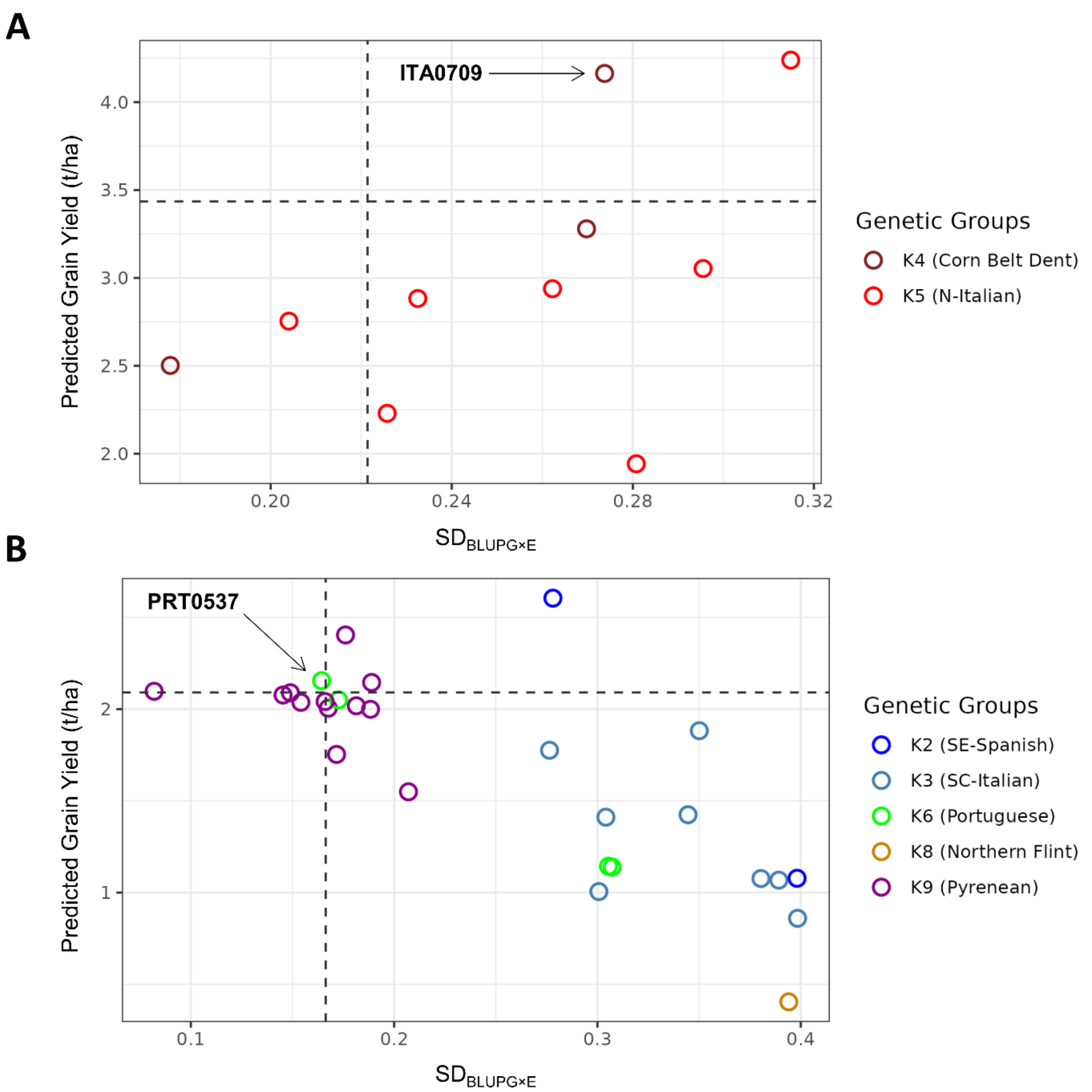

**Figure S14: Selection of the highest performing and most stable landraces with low genomic offset in NE-Romania (A) and SW-France (B) for future climatic scenario SSP585 and time horizon 2081-2100.** Relationship between predicted grain yield and stability for landraces with low genomic offset (GO <0.03) for target regions of NE-Romania (**A**) and SW-France (**B**). Grain yield is predicted by the genomic prediction model M4 that accounts for GO and within-population gene diversity (Hs). Grain yield stability is estimated with the standard deviation of genotype-by-environment best linear unbiased predictors ($SD_{BLUPGxE}$). Arrows highlighted two promising non-local landraces, that combine relative high yield and stability, for target regions and that could be considered as interesting genetic resources for adapting maize to this future climatic scenario.

#### Figure S15 – Prediction of Yield Change based on Genomic Offset effect and Genetic Offset predictions for past and future (2081-2100 SSP585) climate

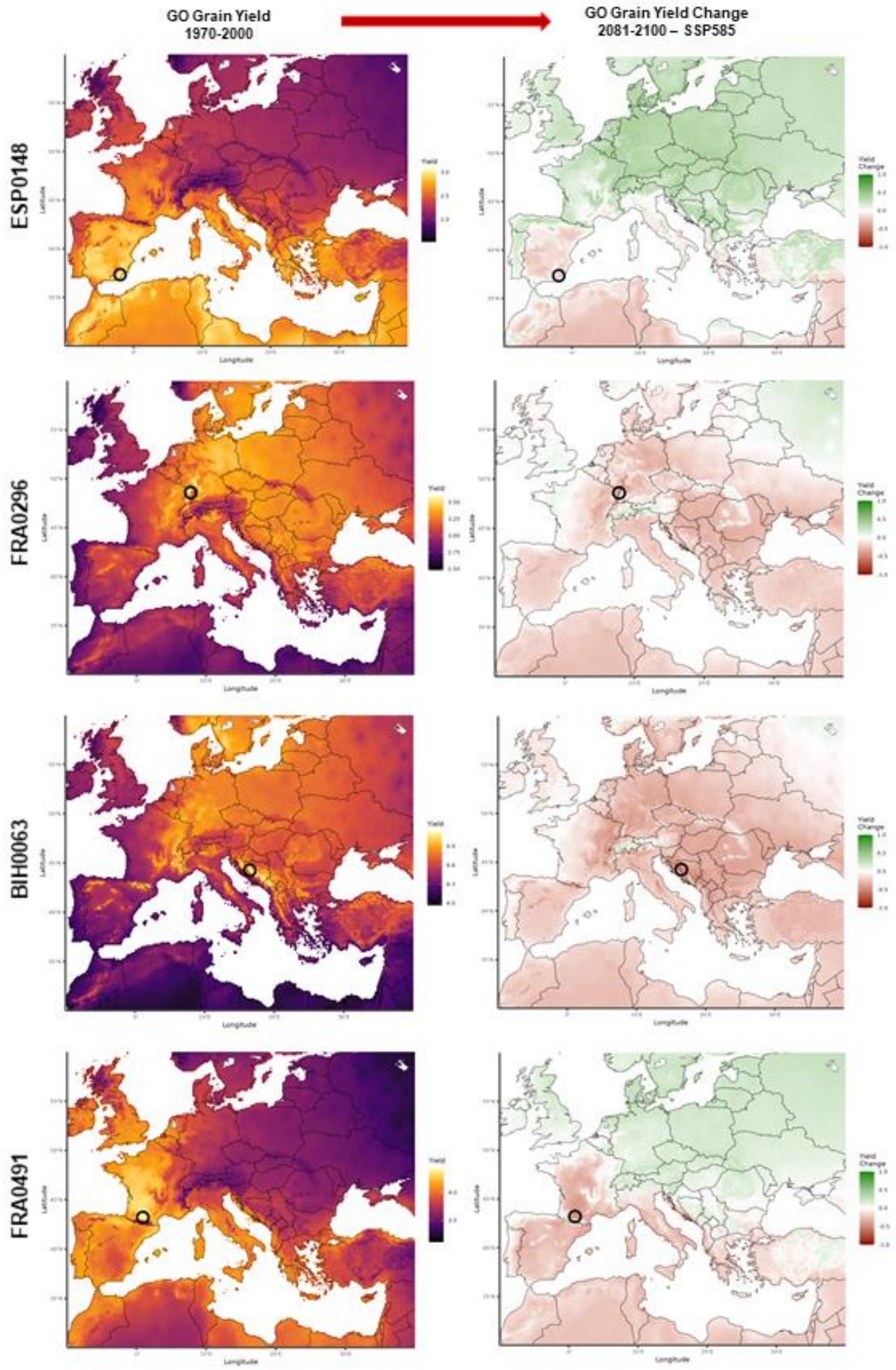

**Figure S15: Yield variation and change due to genomic offset across Europe for period 2081-2100 under SSP585 scenario.** Maps representing yield variation due to genomic offset in past climatic conditions (left) and yield change due to genomic offset under future scenario conditions (right) for ESP0148, FRA0296, BIH0063, and FRA0491. Future scenario maps show that some landraces are expected to benefit (regions in green) or suffer (regions in red) in predicted future climatic conditions as yield due to genomic offset would decrease or in-crease, respectively.
